# The mouse gut microbiota responds to predator odor and predicts host behavior

**DOI:** 10.1101/2025.07.01.662568

**Authors:** Madalena V. F. Real, Maren N. Vitousek, Michael J. Sheehan, Andrew H. Moeller

## Abstract

Chronic stressors can alter the mammalian gut microbiota in ways that mediate host stress responses, but the impacts of acute stressors on these interactions are less well understood. Here, we show that brief exposure of wild-derived mice to predator odor altered gut-microbiota composition, which in turn predicted host behavior. We investigated the individual and combined effects of 15-minute exposures to synthetic fox fecal odor and 30 days of chronic social isolation, an established chronic stressor. Using ethological assays, visceral adipose tissue transcriptomics, and genome-resolved metagenomics, we found that predator-odor exposure significantly affected mouse behavior, gene expression, and gut microbiota. Predator odor–responsive bacteria were associated with the expression of genes involved in anti-microbial defense, and host behavioral responses were predicted by random forest models trained on gut-microbiota profiles. These findings indicate interactions between the gut microbiota and wild-mouse responses to the threat of predation, an ecologically relevant acute stressor.

## Introduction

Animals are faced with diverse stressors that vary in duration and intensity^1,2^, giving rise to distinct physiological and behavioral responses^3,4^. In mammals, behavioral^5–7^, endocrine^8,9^, and immune^5,10,11^ stress responses are mediated in part by the community of host-associated microorganisms that reside in the gastrointestinal tract, i.e., the gut microbiota^12^, which is shaped by changes in the internal and external host environment^13–15^. Recent work has shown that chronic stressors, such as social isolation, can alter the gut microbiota in ways that mediate host stress responses^16–20^. In contrast, the effects of acute stressors on the gut microbiota, and the relationships between the gut microbiota and host responses to acute stress, remain less well-understood.

In rodents, the threat of predation represents an intense acute stressor, and animals have evolved to use environmental cues, such as predator-odor marks, to adjust their behavior and avoid predator encounters^21^. Previous work in mice has shown that a single brief exposure to synthetic fox fecal odor 2,5-dihydro-2,4,5-trimethylthiazoline (TMT) can lead to long-lasting behavioral changes, such as increased fearfulness^22–26^. However, the effects of acute exposure to predator odor on the gut microbiota, and the extent to which predator odor–induced changes in the gut microbiota interact with host stress responses, have yet to be investigated.

Here, we tested the effects of acute exposure to predator odor on the gut microbiota, gene expression, and behavior of wild-derived house mice (*Mus musculus domesticus*) and contrasted these effects with those of social isolation, a well-established chronic stress paradigm. We longitudinally sampled the gut microbiota of males and females exposed to predator odor for 15 minutes at the beginning of the third and fourth weeks of the experiment or socially isolated for 4 weeks. These wild-derived mice retain a more diverse gut microbiota than do lab mice and have recently been proposed as a solution to the microbiota-related issues around the poor reproducibility and low translational value of laboratory mouse studies^27,28^. We found that brief exposure to predator odor had a greater impact on the gut microbiota than did prolonged social isolation. Additionally, the taxonomic composition of the gut microbiota, but not host gene expression profiles, significantly predicted inter-individual variation in host behavior. These findings support links between the gut microbiota and host responses to acute stressors.

## Results

### Predator-odor exposure and social isolation gave rise to stress-associated behaviors

We compared the effects of brief exposure to predator odor to those of prolonged exposure to social isolation—a well-established chronic stress model. All experiments were conducted using wild-derived mice originally captured in Saratoga Springs (New York, USA) in 2012 and subsequently inbred in the lab^29–31^. At the beginning of the experiment (D0), we separated groups of 6-week-old same-sex littermates into pair- and single-housed mice (Figure 1A). Two weeks later (D15), we exposed these mice to 35 µL of either synthetic fox fecal odor (2,5-dihydro-2,4,5-trimethylthiazoline, TMT) or a control scent (deionized water, H_2_O) for 15 minutes. We repeated this exposure procedure the following week (D22). The full factorial design produced 9 unstressed control cages (10 male and 8

**Figure 1.**
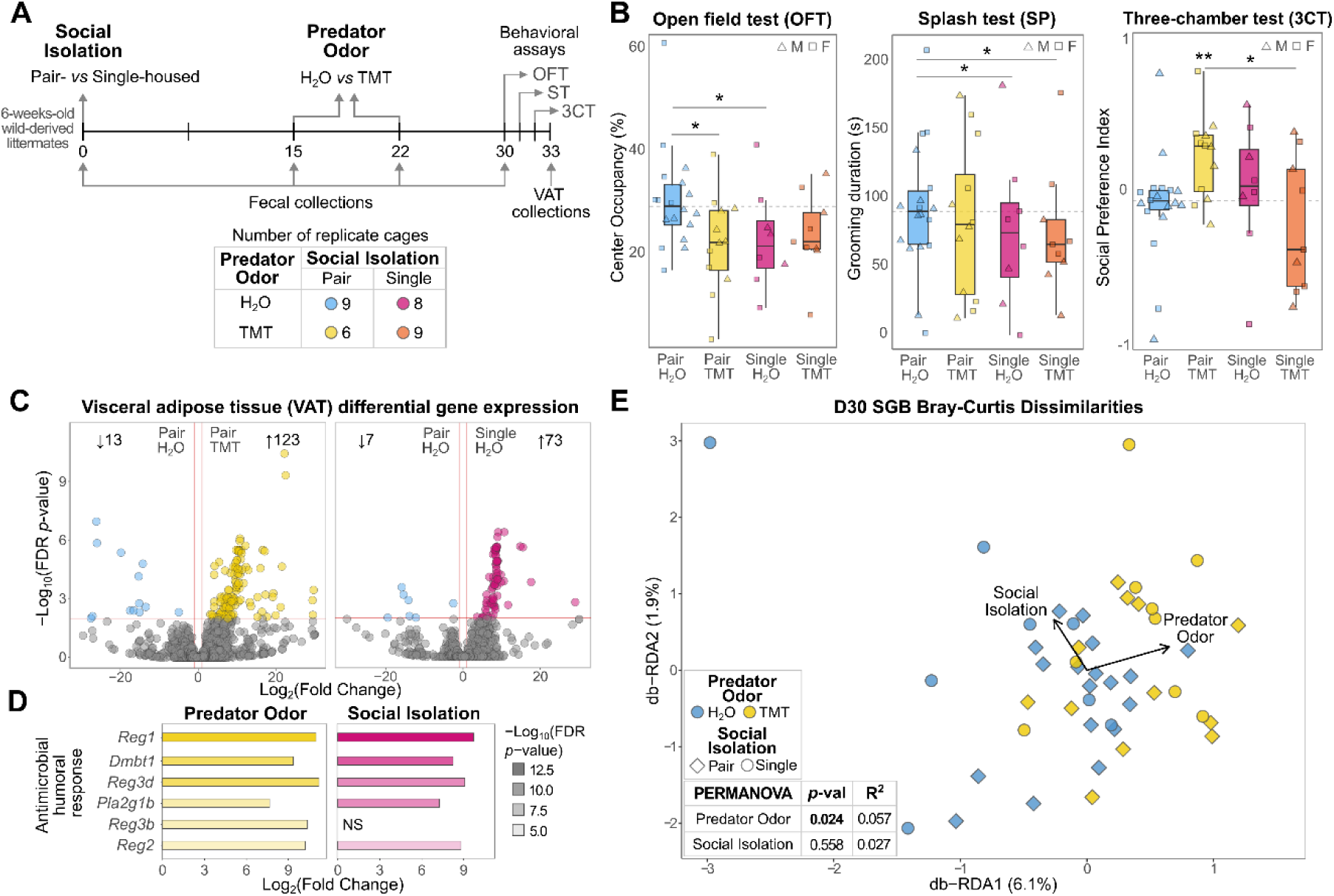
Predator-odor exposure altered host behavior, gene expression, and gut microbiota. (**A**) Diagram shows experiment timeline and number of cages per treatment group. (**B**) Box plots show, for male (triangles) and female (squares) mice, time spent in the central area of open-field tests (OFT), time spent grooming in splash tests (SP), and social preference index in three-chamber tests (3CT). Differences between stressor groups and unstressed controls were tested using linear mixed-effects models (LMM) accounting for sex, litter, and cage effects (* = *p* < 0.05). Differences between stressor groups were assessed by pairwise comparisons of LMM marginal means (* = FDR-adjusted *p* < 0.05). In the 3CT plot, we tested whether the social preference index within each group significantly differed from zero with a one-sample Wilcoxon test (** = *p* < 0.01). Dashed gray lines indicate unstressed controls’ mean. (**C**) Volcano plots show log2-transformed fold change (LFC) in gene-expression level (points) in the visceral adipose tissue (VAT) of male and female mice in response to predator-odor exposure (left) or social isolation (right). Genes in the upper-outer quadrants were differentially expressed (DEGs; absolute LFC > 1 and FDR-adjusted *p* < 0.01). The total number of DEGs is presented at the top of each plot. (**D**) Bar plots show absolute LFC of genes belonging to the anti-microbial humoral response biological pathway, which was found to be significantly enriched in the up-regulated DEGs of mice exposed to predator odor (left) and social isolation (right). Genes are ordered and colored by the −log_10_(FDR-adjusted *p*) resulting from differential expression analyses. (**E**) Distance-based redundancy analysis (db-RDA) shows the first two axes of constrained ordination of the Bray-Curtis dissimilarity matrix of species genome bins (SGBs) composition at D30, accounting for sex and litter effects. Points represent the SGB profiles of pair (diamond) and single-housed (circle) mice, colored by whether they were exposed to H_2_O (blue) or TMT (yellow). Vectors indicate direction and relative strength of association between stressors and gut microbiome composition. The percent variation explained by each axis is enclosed in parentheses.

female pair-housed H_2_O-exposed mice), 6 predator-odor cages (6 male and 6 female pair-housed TMT-exposed mice), 8 social-isolation cages (3 male and 5 female single-housed H_2_O-exposed mice), and 9 cages that experienced both stressors (4 male and 5 female single-housed TMT-exposed mice; Supplementary Data 1).

We assessed whether brief predator-odor exposure and prolonged social isolation acted as stressors in wild-derived mice by examining the effects of these treatments on host behavior with linear mixed-effects models (LMM) that accounted for sex, litter, and cage effects (Figure 1B; Table S1). We found that individual exposure to the stressors increased mouse fearfulness, as both pair-housed mice exposed to TMT (β = −0.093 ± 0.038, *t*(26.6) = −2.426, *p* = 0.022) and socially isolated mice exposed to H_2_O (β = −0.084 ± 0.041, *t*(39.3) = −2.039, *p* = 0.048) significantly reduced occupancy of the exposed central area during the open field test ^32^ when compared to unstressed controls. Mice that experienced both stressors displayed a non-significant trend toward increased fearfulness (β = −0.072 ± 0.039, *t*(39.9) = −1.831, *p* = 0.075). We conducted additional pairwise comparisons between the stressor groups but found no significant differences in center occupancy (Table S2). There were also no significant differences in overall locomotion between the treatment groups (Table S1 and S2), indicating that the observed reduction in center occupancy was not due to a decrease in overall movement. These results confirm that, in line with previous work in lab mouse strains^3,16,33^, predator-odor exposure and social isolation functioned as stressors that increased fear-associated behaviors in wild-derived mice.

Predator-odor exposure has also been shown to lead to a reduction in self-grooming^25^, which is part of a suite of behaviors that are inhibited when rodents enter a more vigilant state after the threat of predation^34^. We assessed grooming behavior with the splash test^35^, where individual mice were sprayed with a palatable sucrose solution and their grooming behavior measured over the subsequent 5 minutes. Social isolation, but not predator-odor exposure, significantly decreased grooming behavior, such that both socially isolated H_2_O-exposed mice (β = −44.327 ± 20.299, *t*(42.0) = −2.184, *p* = 0.035) and mice that experienced the combined stressors (β = −39.629 ± 19.204, *t*(41.9) = −2.064, *p* = 0.045) spent significantly less time grooming than did unstressed controls (Figure 1B; Table S1). There were no significant differences in grooming behavior between stressor treatments (Table S2).

Both brief predator-odor exposure^36^ and prolonged social isolation^16,33^ have been previously reported to decrease rodent sociability, here measured by the relative time spent interacting with a social stimulus (stranger) *versus* a non-social stimulus (empty cup) in the three-chamber test^37^. To assess the sociability of each treatment group, we compared the time each mouse spent interacting with either the social or non-social stimulus using a paired Wilcoxon test. We complemented these analyses with a one-sample Wilcoxon signed-rank test to assess whether the average social preference in each treatment group was significantly different than zero. Finally, we investigated whether the social preference index of mice that experienced the stressors was significantly different than that of controls with a LMM. Contrary to prior work in lab mice^36^, we found that pair-housed wild-derived mice exposed to predator odor displayed increased sociability, spending significantly more time interacting with the social stimulus than the non-social stimulus (paired Wilcoxon: *V* = 75, *p* = 0.002; Figure S1) and displaying a social preference index significantly greater than zero (one-sample Wilcoxon: *V* = 74, *p* = 0.003; Figure 1B). In contrast, the social preference index of both groups of single-housed mice or unstressed controls did not differ significantly from zero. While we found no significant differences in the social preference index of mice exposed to the stressors compared to controls (Table S1), the social preference of pair-housed mice exposed to TMT was significantly higher than that of their single-housed counterparts (β = 0.513 ± 0.197, *t* = 2.610, FDR-adjusted *p* = 0.027). We did not observe significant sex differences in any of the assayed behaviors (Table S1). Overall, our results show that both brief exposure to predator odor and prolonged social isolation resulted in an increase in stress-related behaviors of wild-derived mice (i.e., fearfulness and, in the case of socially isolated mice, reduced self-grooming), and that predator-odor exposure in the context of pair housing increased sociability.

### Predator-odor exposure altered the transcriptome of an immunological tissue

Given the observed effects of brief exposure to predator odor and prolonged social isolation on the behaviors of wild-derived mice, we next investigated the impacts of these stressors on gene expression profiles in the visceral adipose tissue (VAT). Both acute and chronic stressors have been shown to increase gut permeability, leading to the translocation of microbes and their metabolites from the gut lumen to surrounding tissues^38^, such as the VAT. As an immunologically active organ^39^, the VAT can detect these microbial signals^40^ and inhibit bacterial infection and sepsis^41,42^ through the secretion of anti-microbial peptides (AMPs)^43,44^. To investigate possible effects of the stressors on gene expression profiles, we used 3’ RNA-seq to sequence the transcriptomes of VAT collected at the end of the experiment (D33). A total of 35 samples met our quality threshold of over 1 million transcripts/sample, averaging 6.54 million (SD = 3.22 million) transcripts/sample (Supplementary Data 2).

We found that TMT exposure, but not social isolation, significantly altered the VAT gene expression profile (PERMANOVA of Euclidean distances accounting for sex and litter; predator odor: R^2^ = 0.047, *F*(1) = 2.185, *p* = 0.035; social isolation: R^2^ = 0.037, *F*(1) = 1.681, *p* = 0.097; Figure S2; Table S3). We tested for differentially expressed genes (DEGs; absolute log_2_(fold change) (LFC) > 1 and FDR-adjusted *p* < 0.01; Supplementary Data 3) between mice exposed to individual stressors and unstressed controls. While there was considerable overlap in DEGs between the different stressor treatments (Figure S3), we found that, compared to social isolation, predator-odor exposure led to a greater number of DEGs (Figures 1C and S3) and a significantly larger mean fold-change in gene expression (unpaired Wilcoxon test *p* = 0.026; Figure S4). Interestingly, DEGs up-regulated in both TMT-exposed (FDR-adjusted *p* = 0.007) and socially isolated mice (FDR-adjusted *p* = 0.004) were significantly enriched in biological pathways related to anti-microbial humoral immune responses, including those mediated by anti-microbial peptides (AMPs; Figure 1D; Table S4). These results show that while both stressors activated anti-microbial defenses, brief predator-odor exposure had a greater overall impact on VAT gene expression than did prolonged social isolation.

### Predator-odor exposure had a greater impact on the gut microbiota than did social isolation

We assessed the impacts of brief exposure to predator odor and prolonged social isolation on the gut microbiota by sequencing the metagenomes of fecal samples collected at the beginning (D0) of the experiment, immediately before the TMT exposures (D15 and D22), and the end of the experiment (D30; Figure 1A; Supplementary Data 4A). Sequencing D0 and D30 samples yielded a mean of 33.6 million (SD = 5.0 million) reads/sample. Intermediate samples from D15 and D22 were sequenced at lower depth, yielding a mean of 10.0 million (SD = 2.4 million) reads/sample. We assembled and binned D0 and D30 reads into metagenome-assembled genomes (MAGs) using a custom automated workflow (MAGmaker), following previously published methods^45^. The MAGmaker workflow yielded 6,404 MAGs, 3,542 of which were high quality (>90% completeness, <5% contamination). We then created a custom database of 173 representative species-level genome bins (SGBs) by dereplicating high-quality MAGs at 95% average nucleotide identity (ANI). SGBs were classified with the Genome Taxonomy Database^46,47^ (Supplementary Data 4B), and their relative abundances in all samples were quantified using InStrain^48^ (Supplementary Data 4C). The mean mappability of our samples to the SGBs was 75.75% (SD = 3.04%), indicating that the custom-made SGB database was generally representative of the bacterial diversity in our samples. Samples were rarefied to the minimum library size before beta-diversity analyses (8,470,661 reads for D0 and D30 samples, and 3,896,354 reads for D15 and D22 samples; Supplementary Data 4D).

We tested the effects of predator-odor exposure and social isolation on the SGB profile through PERMANOVA analyses of Bray-Curtis dissimilarities, accounting for sex, litter, and cage effects. While social isolation significantly affected the mean SGB composition at D22 (PERMANOVA R^2^ = 0.029, *F*(1) = 2.546, FDR-adjusted *p* = 0.008), this effect disappeared a week later (PERMANOVA R^2^ = 0.011, *F*(1) = 1.282, FDR-adjusted *p* = 0.277; Figure S5). Instead, at D30, social isolation tended to increase inter-host variance in microbiota composition (PERMDISP *F*(1) = 5.754, FDR-adjusted *p* = 0.076). In contrast, TMT exposure impacted the gut microbiota not by changing its inter-individual variance (PERMDISP *F*(1) = 0.006, FDR-adjusted *p* = 0.936) but by significantly shifting the mean SGB composition compared to H_2_O exposure (PERMANOVA R^2^ = 0.026, *F*(1) = 2.930, FDR-adjusted *p* = 0.008). No such effects were observed before predator-odor exposure at D0 and D15 (Table S5).

To further examine the effects of stressor exposure while controlling for potential confounding variables, we performed a distance-based redundancy analysis (db-RDA), a constrained ordination approach that tests whether variation in gut-community composition can be explained by predator-odor exposure or social isolation after accounting for sex and litter effects. We found that TMT exposure (PERMANOVA partial R^2^ = 0.057, *F*(1) = 1.992, *p* = 0.024), but not single housing (PERMANOVA partial R^2^ = 0.027, *F*(1) = 0.900, *p* = 0.558), drove a significant shift in the SGB profile of mice sampled one week after the second TMT exposure (D30; Figure 1E). Overall, these results indicate that two acute exposures to TMT were sufficient to shift the SGB profiles of the mouse gut microbiota, and that by the end of the experiment, the impacts of brief predator-odor exposure on gut-microbiota composition were greater than those of prolonged social isolation.

### Predator-odor exposure drove consistent alterations in the relative abundances of specific SGBs

Given the observed changes in gut-microbiota composition following predator-odor exposure, we investigated which specific microbial taxa drove these patterns. We found that a single exposure to TMT was sufficient to significantly alter the relative abundances of several SGBs (D22 absolute LFC > 1 and FDR-adjusted *p* < 0.01; Figure 2A; Supplementary Data 5). Seven of these SGBs remained differentially abundant (DA) one week after the second TMT exposure (D30), with the majority being enriched in TMT-exposed mice and only one in H_2_O-exposed mice (Figures 2A and 2B). In contrast, seven out of the eight SGBs that responded consistently to social isolation were enriched in pair-housed mice, with only one SGB enriched in single-housed individuals (Figures 2A and 2B). The latter, SGB158, was the only species consistently enriched in both stressor groups, suggesting a shared response, while most other consistently responsive SGBs showed stressor-specific patterns of enrichment. These findings indicate that a subset of SGBs in the gut microbiota of wild-derived mice displayed consistent changes in abundance in response to brief predator-odor exposure and prolonged social isolation across two timepoints (D22 and D30).

**Figure 2.**
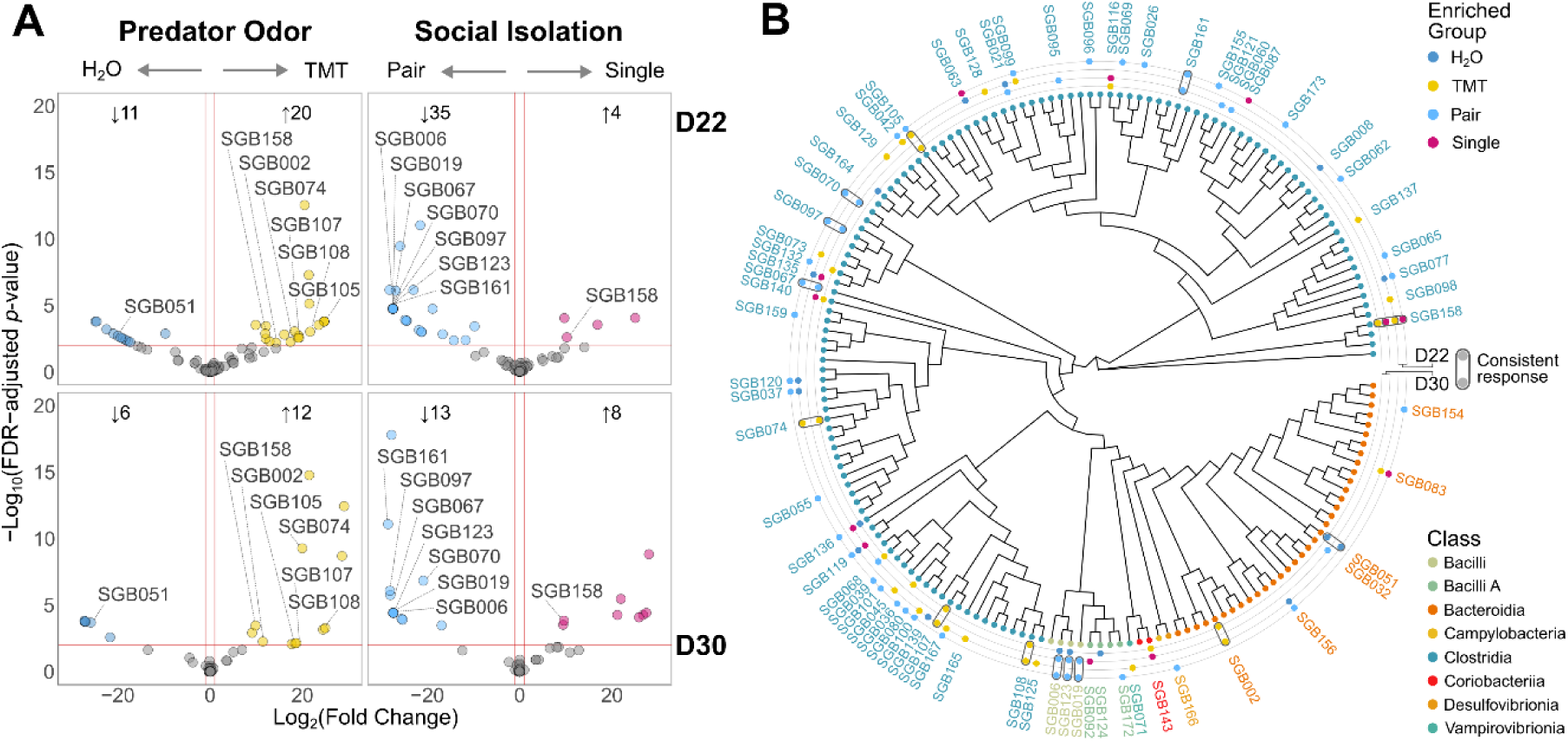
Predator-odor exposure drove consistent alterations in the relative abundances of specific SGBs. (**A**) Volcano plots show log_2_-transformed fold change (LFC) in the relative abundances of species genome bins (SGBs; points) in the gut microbiota in response to predator-odor exposure (left column) or social isolation (right column) a week after the first TMT exposure (D22; top row) and a week after the second TMT exposure (D30; bottom row). Upper-outer quadrants contain points corresponding to differentially abundant SGBs (DESeq2; absolute LFC > 1 and FDR-adjusted *p* < 0.01). The total number of differentially abundant SGBs is presented at the top of each plot. SGBs that responded consistently to the stressors at both D22 and D30 are labelled. (**B**) Phylogenetic tree shows evolutionary relationship (number of amino acid substitutions) between SGBs, with the tips colored by bacterial class. The rings show differentially abundant SGBs at D22 (outer two rings) and D30 (inner two rings), colored by the stressor group in which they were enriched: H_2_O (dark blue) *vs.* TMT (yellow) and Pair (light blue) *vs.* Single (pink). SGBs that displayed a consistent response to the stressors across both D22 and D30 are encircled.

### Stressor–responsive SGBs co-varied with host anti-microbial and behavioral responses

As the stressors up-regulated the expression of genes involved in AMP production, we investigated possible associations between the expression of these AMP-associated genes and the relative abundances of bacteria that responded consistently to predator-odor exposure or social isolation. We conducted these analyses both across and within treatment groups. In the within-group analyses, we controlled for the effects of the stressor treatments by testing associations with partial Spearman correlations, that residualized host phenotypes (i.e., AMPs expression levels and behavior) and SGB relative abundances against each treatment group. We found that, a week after the first predator-odor exposure (D22), TMT-responsive SGBs generally showed positive associations with genes involved in AMP production, although, after correcting for multiple testing, this correlation was only significant for SGB074—identified as Oscillospiraceae species *Pelethomonas* sp. 009774055 (Figure 3). Interestingly, TMT-responsive SGB105—identified as Lachnospiraceae species *Acetatifactor* sp. 011959105—was negatively associated with all genes involved in AMP production. This inverse relation with anti-microbial immune responses was also found in SGBs depleted in mice exposed to the stressors, such SGB051—identified as Muribaculaceae species *Lepagella* sp. 002361155 and enriched in H_2_O-exposed mice—as well as most SGBs that consistently responded to pair-housing (Figure S6). These results show that the relative abundances of stressor–responsive SGBs were significantly associated with host anti-microbial defenses.

**Figure 3.**
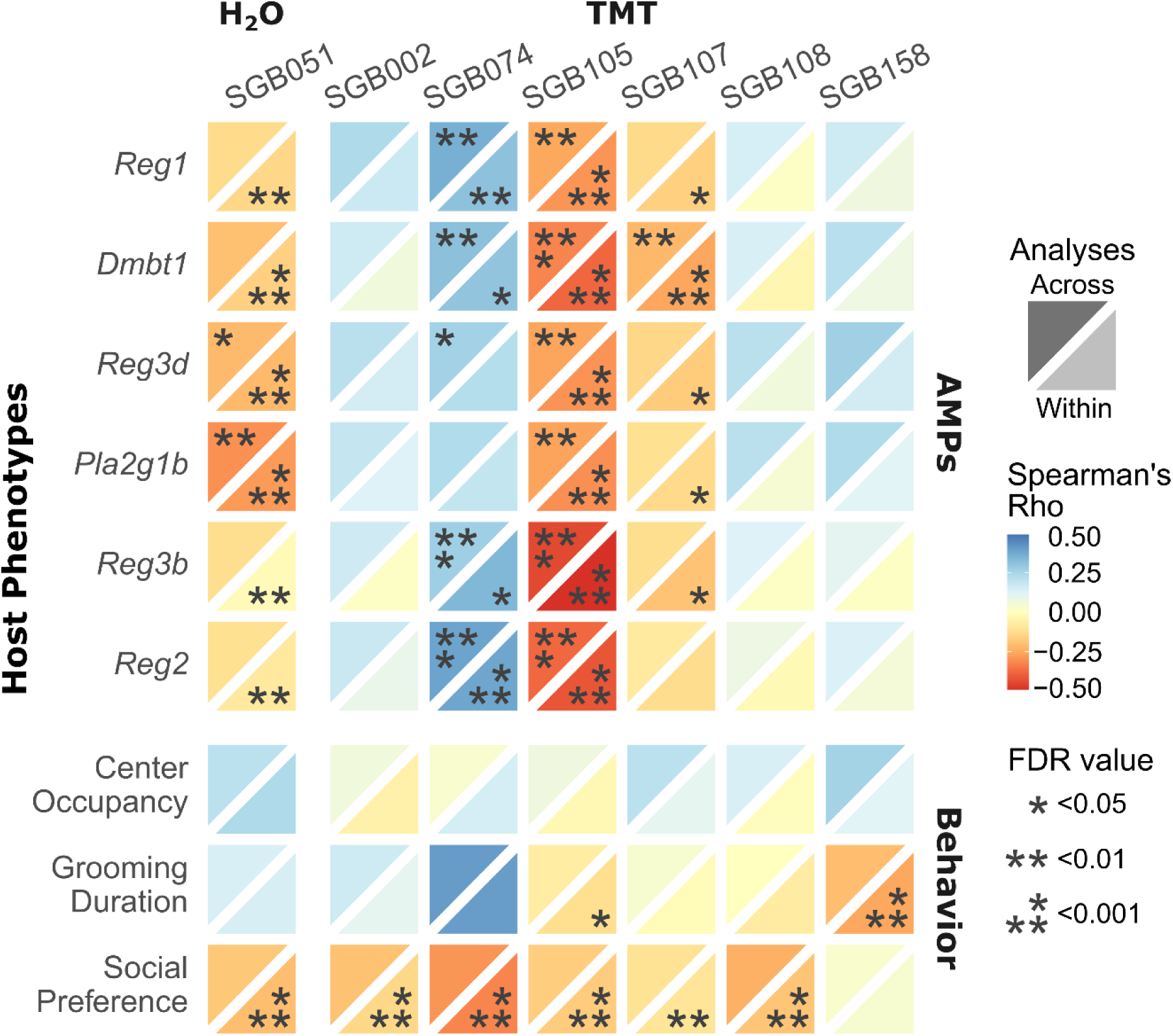
Predator odor–responsive SGBs co-varied with host anti-microbial and behavioral responses. Heatmap shows associations between the D22 relative abundances of SGBs that were consistently associated with predator-odor exposure (differentially abundant at both D22 and D30) and host anti-microbial (AMPs) and behavioral responses. The top facet represents correlations across treatment groups, and the bottom facet correlations within groups. Facets are colored according to Spearman’s Rho and annotated with the correlation’s FDR-adjusted *p*-value (* < 0.05; ** < 0.01; *** < 0.001).

We also tested whether the relative abundances of stressor–responsive SGBs co-varied with host behavioral responses. We found that within-group variation in D22 relative abundances of all stressor–responsive SGBs except SGB158 were negatively associated with mouse social preference (Figures 3 and S6). Instead, this Lachnospiraceae species, enriched in both TMT-exposed and socially isolated mice, was negatively correlated with grooming behavior. These findings indicate that behavioral responses to stressor exposure, such as reduced grooming and social preference (particularly for single-housed mice exposed to TMT), could be partially explained by the relative abundances of stressor-responsive SGBs measured a week prior.

### Gut microbiota predicted host behavioral responses to stressors

Motivated by the observation that host phenotypes could be partially explained by the relative abundances of specific stressor-responsive SGBs, we tested the predictive power of the entire gut microbiota. Variation in gut microbial communities has been shown to shape host behavioral responses to stress^5–7^. Here, we aimed not to infer causality directly but to assess whether inter-individual differences in gut microbiota composition contain information that can be used to predict variation in host behavior both across and within treatment groups. To this end, we trained random forest models to predict host behavior based on SGB relative abundances at each timepoint. For within-group analyses, the predicted behavioral variables were residualized against stressor treatment. The model’s performance was assessed through a 5-fold cross-validation strategy, and we conducted a permutation test to measure the statistical significance of the observed R^2^ values, which was corrected to account for multiple comparisons across predicted variables and the four timepoints (Table S6). Additionally, we used linear regressions to test the relation between the observed behavioral values and the ones predicted by our models. We found that by the end of our experiment (D30), the gut microbiota consistently predicted grooming and social behavior both across and within treatment groups (Figure 4A). Grooming and fearfulness could also be predicted by inter-individual variation in SGB relative abundances sampled a week prior (D22; Figure S7A). In contrast, the gut microbiota sampled before TMT exposure, at either D0 or D15, was not predictive of host behavior. Interestingly, all SGBs found to respond consistently to predator-odor exposure ranked among the top quarter most important features for predicting at least one of the behavioral phenotypes (Figure 4B; Table S7). For example, the D22 abundance of the predator–odor and social–isolation-responsive SGB158 was the 3^rd^ most important feature for predicting grooming behavior across treatments more than a week later (Figure S7B), corroborating our Spearman correlation results. Furthermore, we found that the gut microbiota was a better predictor of host behavior than was VAT gene expression, which did not significantly predict behavioral phenotypes (Figure S8; Table S6). These results indicate that the gut microbiota sampled at the end of the experiment, a week after the second TMT exposure, significantly predicted subsequent host behavioral phenotypes.

**Figure 4.**
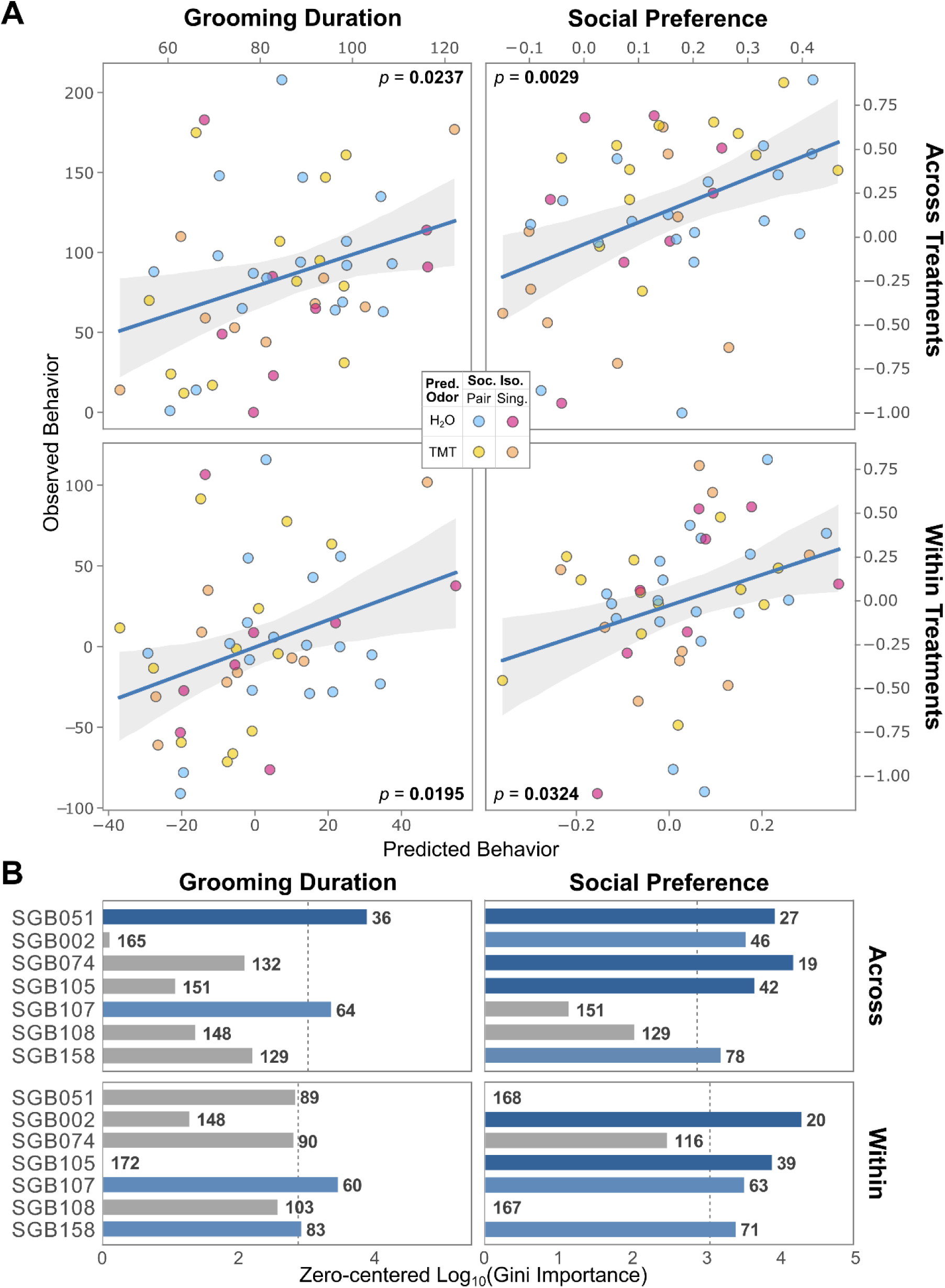
Gut microbiota predicted host behavior. (**A**) Scatter plots show the relationships between observed grooming (left column) and social (right column) behavior and the values predicted by a random forest model trained on SGB relative abundances at D30, either across (top row) or within treatment groups (bottom row; behavior residualized to account for the effect of stressor treatment). Points are colored by treatment group: blue, Pair H_2_O; yellow, Pair TMT; pink, Single H_2_O; orange, Single TMT. The significance of the linear regression is shown in the upper inner corner, with significant values (*p* < 0.05) in bold. (**B**) Bar plots show the zero-centered log_10_ (Gini Importance) of SGBs consistently associated with predator-odor exposure (differentially abundant at both D22 and D30) for predicting grooming (left) and social (right) behaviors both across (top) and within (bottom) treatment groups. The SGB importance rank order is annotated in bold. The bars are colored light blue if the SGB was among the top half most important features and dark blue if among the top quarter. The dashed line indicates the median importance value.

## Discussion

We found that acute exposure of wild-derived mice to predator odor promoted stress-associated behaviors, altered gene expression patterns in the visceral adipose tissue through the activation of the AMP-mediated anti-microbial responses, and shifted the species composition of the gut microbiota. Surprisingly, we found that brief exposure to predator odor had a larger impact on the VAT transcriptome and the gut SGB profile than did prolonged social isolation—a well-established chronic stress model previously shown to impact the gut microbiota^16,17^. Moreover, variation in gut-microbiota composition at the end of the experiment (D30) predicted host behaviors. These results show that predator odor alters the murine gut microbiota and suggest links between inter-individual variation in the gut microbiota and behavioral responses in the context of predation stress.

By comparing the effects of two distinct stressors and their combination on the gut microbiota, host gene expression, and behavior, our study builds on prior work focused on the effects of individual stressors^16,17,49,50^. We observed that the combination of stressors often had non-additive effects, exemplifying the context dependence of the stress response^3^. For example, we observed that pair-housed mice exposed to predator odor displayed significantly higher social preference than did their socially isolated counterparts, demonstrating that the social environment can affect the response to consequent stressors. In our case, pair-housed mice might seek out proximity with conspecifics when under the threat of predation—a behavior described as huddling^51^— while single-housed mice exposed to predator odor display the asocial behavior commonly observed after chronic stressor exposure^52^.

Including four treatment groups to test the effects of two stressors and their combination constrained sample sizes. This limitation may have reduced our power to detect, for example, a significant decrease in center occupancy for mice that experienced both stressors compared to controls, or a significant effect of social isolation on gut microbiota composition, as has been previously reported^16,17^. Nevertheless, our group sizes offered sufficient statistical power to detect multiple significant effects of the stressors on mouse behavior, VAT gene expression, and gut microbiota. The biological significance of these results is further supported by the inclusion of both male and female mice in our experimental design, thereby addressing potential sex biases in the stress response^53–55^.

We identified both general and stressor-specific responses in wild-derived mice and their gut microbiota. For example, we observed that both stressors up-regulated AMP expression in the VAT. Interestingly, the expression of these AMPs co-varied with the relative abundances of stressor-responsive taxa, even after accounting for stressor effects (Figure 3). For example, SGB051, which was depleted in TMT-exposed mice, was negatively associated with all genes involved in AMP production. This SGB was identified as a Muribaculaceae, a family previously associated with anti-inflammatory markers^56,57^ and often depleted in inflammatory bowel disease (IBD)^56,58^. Muribaculaceae produce short-chain fatty acids (SCFAs), which have been shown to stimulate mucus production and reduce inflammation^59^. The associations observed in our study could be due to either SGB051 growing better in the absence of AMPs or to a negative effect of this species on anti-microbial responses. One possible explanation for our results is that gut-microbiota responses to predator odor were mediated by VAT inflammation triggered during the acute stress response. Alternatively, it is also possible that predator-odor exposure led to changes in the gut microbiota that triggered VAT inflammation. Given that both increased VAT inflammation^60^ and stress-induced changes in the gut microbiota^16,17,61^ have been shown to affect mouse behavior, the observed behavioral effects of stressor exposure may be mediated by signaling along the gut-brain-immune axis^62–64^. Alternatively, the gut microbiota may interact with host behavior in a VAT-independent manner, as variation in gut species composition was a better predictor of mouse fearfulness, grooming, and sociability than was the VAT gene expression profile. Understanding the mechanisms underlying the associations between the predation stress–altered microbiota, host inflammatory responses, and behavior will require controlled experiments in germ-free animals designed to test the effects of stressor-associated microbiota on mouse phenotypes that can complement previous work showing that host responses to acute stressors are microbiota-dependent^65–67^.

Several results contradicted expectations based on previous literature using lab mouse strains. For example, while four weeks of social isolation was sufficient to alter the gut microbiota of C57BL6 males^16,17^, in our study, only predator-odor exposure significantly shifted the mean species composition of the gut microbiota at D30. This discrepancy could be due to differences in the robustness of the gut community to perturbation between mouse strains. Lab mice have a depleted, and potentially more volatile, gut microbiota compared to wild mice^27,28^, and previous work has shown that wild-derived mouse lines can retain microbiota from wild populations for over a dozen generations in the lab^68^. Additionally, there is evidence that wild-derived mice are less sociable than lab mouse strains^69–71^, potentially increasing their tolerance for social isolation. This hypothesis is supported by our observation that unstressed controls lack a clear social preference, whereas commonly used lab lines tend to display high sociability^72^. The sociability displayed by pair-housed wild-derived mice exposed to predator odor similarly contradicts findings of previous studies of lab mice^36^. Brief predator-odor exposure is likely a more potent stressor for wild-derived mice than prolonged social isolation. This is supported by previous work showing wild-derived mice are more readily responsive to the threat of predation than lab mice^73^. Thus, this work adds to a growing body of evidence motivating the incorporation of wild-derived mice into research as a biologically relevant complement to the work conducted on classic lab strains^27,28,69,74^. Moreover, our results show that acute stressors, such as predator-odor exposure, can have effects of comparable or even greater magnitude than a well-established chronic stressor like social isolation.

## Materials and Methods

All procedures conformed to guidelines established by the U.S. National Institutes of Health and have been approved by the Cornell University Institutional Animal Care and Use Committee (protocol #2015-0060).

### Animals

We used a wild-derived *M. m. domesticus* line, NY3, which was captured in Saratoga Springs (NY, USA) in 2012 and subsequently inbred in captivity^29–31^. These mice are related to the SAR/NachJ lines currently provided by the Jackson Laboratory^74^. Mice were kept on an inverted light cycle (light:10pm-10am) and all handling was conducted during the active (dark) phase using red light. Standard chow diet and water were available *ad libitum*.

### Stressor Treatments

#### Social Isolation

Mice were weaned at 3 weeks of age (M = 22 days old, SD = 2) and co-housed with same-sex littermates. At 6 weeks of age (D0; M = 37 days old, SD = 8), we separated co-housed same-sex littermates into pair- and single-housed mice, in a total of 8 male and 7 female mouse pairs, and 7 male and 10 female mice that were socially isolated.

#### Predator-Odor Exposure

Two weeks after the start of the experiment (D15; M = 57 days old, SD = 5), we exposed mice to synthetic fox fecal odor, 2,5-dihydro-2,4,5-trimethylthiazoline (TMT) (90% purity; BioSRQ, Sarasota, FL, USA). Mice were individually placed in an empty cage, brought to a fume hood, and a filter paper with 35 µL of either TMT or deionized water (H_2_O) was placed on their cage metal grid topper for 15 minutes. After the exposure period, we removed the filter paper, keeping the mice in the fume hood for an additional 15 minutes to ensure the TMT was removed from the work area. Afterwards, all mice were returned to their home cages. We exposed control mice first and the TMT-treated mice last, to ensure controls never encountered the predator odor. We repeated this exposure procedure the following week (D22; M = 64 days old, SD = 5). In total, 10 males and 11 females were exposed to TMT, while 13 males and 13 females were exposed to H_2_O. Considering both stressors, there were 9 pair-housed H_2_O-exposed cages (5 male pairs and 4 female pairs), 6 pair-housed TMT-exposed cages (3 male pairs and 3 female pairs), 8 single-housed H_2_O-exposed cages (3 males and 5 females), and 9 single-housed TMT-exposed cages (4 males and 5 females).

#### Behavioral Assays

We assayed mouse behavior on the fifth week of the experiment (D30; M = 72 days old, SD = 6), over three consecutive days. The tests were ordered from least to most disruptive, starting with the Open Field Test (OFT)^32^, followed by the Splash Test (SP)^35^, and ending with the Three-Chamber Test (3CT). All behavior trials were conducted between 1:00-6:00 pm, and as mice were kept on an inverted light cycle, this time corresponded to the animals’ active (dark) phase. The OFT and 3CT were conducted inside a soundproof chamber in a separate behavior room. The mice were habituated to the behavior room for at least 30 minutes. We alternated the testing order between experimental groups and wiped the behavioral apparatuses between trials with 70% ethanol. An infrared camera placed above the apparatuses recorded the behavior tests. All analyses were conducted by the same experimenter, who was blind to the experimental condition.

#### Open Field Test

Mice were placed in the center of a white PVC arena (48 × 46 × 44 cm) and allowed to explore for 10 minutes. We used ToxTrac^75^ to detect mouse trajectories and calculate total distance walked and time spent in the center of the arena (34 × 32 cm; ½ of the area). After the test, each mouse was housed in a new individual home cage.

#### Splash Test

Our protocol was adapted from Nollet ^76^. On the second day of behavior tests, each mouse was placed in an empty “spray” cage and sprayed on their dorsal coat with a 10% sucrose solution. Mice were then quickly returned to their home cage, and grooming behavior was measured for the following 5 minutes. We directly measured the duration of the grooming behavior by using a chronometer to time observed instances of nose/face grooming, head washing, and body grooming, as described by Kalueff and Tuohimaa ^77^.

#### Three-Chamber Test

Mice were placed in the middle chamber of a transparent plexiglass three-chamber arena for a 5-minute habituation period. We then placed overturned wire cups on diagonally opposite corners of each side chamber. One of the cups was empty (non-social stimulus), while the other contained a stranger mouse of the same sex as the test mouse (social stimulus). The position of the stimuli was counterbalanced between trials. We allowed the test mouse to explore the arena for 10 minutes. The recorded video footage was manually scored with BORIS where we logged interactions with the social and non-social stimuli following recommendations by Rein, et al. ^37^. These were behaviors such as directly interacting with the stranger mouse between the cup’s wire bars and/or sniffing the cup or any protruding part of the stranger mouse (e.g., tail). Climbing on top of the cups or grooming in their proximity was not counted as an interaction. Additionally, we calculated the social preference index, which is the difference in interaction time between the social and non-social stimulus, normalized by the total time interacting with both stimuli ^37^. Significantly higher interaction time with the social *versus* non-social stimulus was interpreted as the existence of a social preference^37^.

### Host Gene Expression

#### Tissue Collection and Gross Morphometry

Mice were humanely euthanized with an overdose of CO_2_ followed by cervical dislocation. Immediately after, we dissected the visceral adipose tissue (VAT), which was flash frozen and stored at −80°C.

#### Transcriptomic Sequencing

Total RNA was extracted from the VAT using the RNeasy Lipid Tissue Mini Kit (Qiagen, Germantown, MD, USA). Briefly, approximately 50 mg of VAT were placed in Bead Ruptor 2 mL tubes with 2.4 mm metal beads (Omni International, Kennesaw, GA, USA) and 1 mL of QIAzol Lysis Reagent (Qiagen). The tissues were homogenized with a Bead Ruptor (Omni International) at speed 5 for 30 seconds. We followed the manufacturer’s instructions for the remainder of both RNeasy protocols, including the optional drying step of placing the RNeasy spin column in a new 2 mL collection tube and centrifuging at full speed for 1 minute. Total RNA was eluted twice in 30 µL of RNase-free water (Qiagen). We assessed the concentration and purity of the extracted RNA with a NanoDrop Microvolume Spectrophotometer (Thermo Fisher Scientific) for all samples and with a TapeStation (Agilent Technologies, Santa Clara, CA, USA) for 10% of the samples. The 3’RNA-seq libraries were prepared at Cornell University’s Biotechnology Resource Center (BRC; Ithaca, NY, USA) and sequenced on a NextSeq 500 (Illumina).

#### Transcriptomic Data Processing

We assessed the quality of our transcriptomic data with FastQC (v0.72.0)^78^ and filtered out samples with fewer than 1 million reads. The remaining transcripts were aligned against the *Mus musculus* reference genome (GRCm39 GCF_000001635.27) using STAR (v2.7.10b; setting: --quantMode GeneCounts --outSAMtype BAM SortedByCoordinate)^79^.

### Gut Microbiota

#### Fecal Collections

We sequenced the metagenome from fecal samples collected at the beginning of the first (D0), third (D15), fourth (D22), and fifth (D30) weeks of the experiment. D15 and D22 samples were collected immediately before the predator-odor exposure procedure. D30 samples were collected immediately before the first behavioral assay. All fecal samples were flash frozen and stored at −80°C.

#### DNA Sequencing

Total microbial DNA was extracted with the Quick-DNA MagBead extraction kit (Zymo, Irvine, CA, USA) using an OT-2 liquid handling robot (Opentrons, New York, NY, USA). The metagenomic libraries were prepared with the TruSeq kit (Illumina, San Diego, CA, USA) at Cornell’s BRC. The D0 and D30 libraries were sequenced in a NovaSeq 6000 S4 (PE150 bp; Illumina) at the UC Davis Genome Center (Davis, CA, USA), while the D15 and D22 libraries were sequenced in a NovaSeq X (2 × 150 bp; Illumina) at Cornell’s BRC.

#### Metagenomic Data Processing and Analysis

We processed our metagenomic samples and assembled MAGs using a custom snakemake^80^ workflow (MAGmaker)^45^. Briefly, sequencing adapters were trimmed with Cutadapt (v17.4; setting: “-a AGATCGGAAGAGCACACGTCTGAACTCCAGTCA -A AGATCGGAAGAGCGTCGTGTAGGGAAAGAGTGT -a GGGGGGGGGGGGGGGGGGGG -A GGGGGGGGGGGGGGGGGGGG -u 10 -U 10 --minimum-length 1 --nextseq-trim 20”)^81^, host reads removed with Bowtie2 (v2.5.1; setting: “--very-fast”; host reference genome: GRCm39 GCF_000001635.27)^82^, and the quality of our metagenomic data was assessed with FastQC (v0.72.0)^78^. We assembled high-quality non-host reads from D0 and D30 into contiguous sequences (contigs) using Megahit (v1.2.9)^83^ and assessed assembly quality with Quast (v5.2.0)^84^. We then calculated the distances between our samples using SourMash (v4.4.0)^85^ and selected the 11 samples most dissimilar to all others to be used as “prototypes” against which the metagenomic reads from all samples were mapped with Minimap2 (v2.24)^86^. The resulting coverage information was input into three different binning algorithms: CONCOCT (v0.4.2)^87^, MetaBAT2 (v2.15)^88^, and MaxBin (v2.2)^89^. We then used DASTool (v1.1.6)^90^ to select the optimal set of bins among the three binners. The resulting MAGs were quality controlled for completeness and contamination with CheckM (v1.2.2)^91^ and GUNC (v1.0.6)^92^. We created a custom genome database by dereplicating high-quality MAGs (>90% completeness, <5% contamination) into 95% average nucleotide identity (ANI) species-genome bins (SGBs) using dRep (v3.4.2)^93^. We profiled these SGBs using GTDB-Tk (v2.4.0)^47^. Finally, we calculated the relative abundances of representative SGBs in our samples by mapping all metagenomic reads against our custom genome database with Bowtie2 and profiling them using InStrain (v1.8.0)^48^. The counts from our mapped reads were normalized by the genome size of the representative SGB the read mapped to and the library size of the metagenomic sample from which the read originated. We then converted these normalized counts into a relative abundance percentage. The Bray-Curtis dissimilarities between samples were calculated using methods from the ‘vegan’ package (v2.6)^94^ via ‘phyloseq’ (v1.42.0)^95^.

### Statistical Analyses

All data analyses, except were indicated, were conducted in R (v4.2.2). Means (M) are reported alongside standard deviations (SD). Linear mixed-effects models, controlling for litter and cage as a random effect (formula: Behavior ∼ Sex + Treatment + (1|Litter/Cage_ID)), were implemented via ‘lmerTest’ (v3.1)^96^. Statistical results are shown as estimate (β) ± standard error (SE), followed by *t*-value (with corresponding degrees of freedom), and *p*-value. Pairwise comparisons between stressor treatments were calculated using the ‘multcomp’ package (v1.4-25)^97^. We tested which factors explained the microbiota dissimilarity matrix by running a PERMANOVA ^98^ via adonis2 (formula: Dissimilarity matrix ∼ Sex + Litter + Housing + TMT + Cage) and a PERMDISP^99^ via betadisper, both included in ‘vegan’ (v2.6)^94^. Statistical results are shown in-text as *F* statistic (with corresponding degrees of freedom), R^2^ (applicable to the PERMANOVA, but not PERMDISP), and FDR-adjusted *p*-value to account for the hypothesis testing being repeated across four different timepoints. We calculated normalized counts and differential gene expression or taxa relative abundances between stressor treatments using ‘DESeq2’ (v1.42.1; formula: ∼ Sex + Housing * TMT + Litter)^100^ with LFC shrinkage using the “ashr” adaptive shrinkage estimator^101^, after filtering for genes or taxa with one or more counts in at least three samples. We identified biologically meaningful pathways within DEGs with an enrichment analysis using ‘clusterProfiler’ (v4.10.1)^102^. We investigated whether species relative abundances significantly explained variation in behavior while controlling for stressor treatment effects with a random forest model implemented via Python’s ‘scikit-learn’ (v1.6)^103^. We evaluated our model’s performance using a 5-fold cross-validation strategy, where we subset our data into 5 different “folds” and trained the model 5 times, using 4 of the folds as the training data and the remaining fold as the test data. Our performance metric (R^2^) was averaged across all 5 iterations. This way, each data point is used at least once to both train and test our model, and the R^2^ reflects the model’s performance across the entire dataset. We evaluated the statistical significance of the R^2^ using a permutation test, where we randomized the associations between behavior and species relative abundances and recalculated R^2^ for 1000 iterations, thus generating a distribution of R^2^ values expected under the null hypothesis that behavior is not associated with species relative abundances. We calculated a *p*-value by measuring the likelihood of observing the R^2^ resultant from our random forest in the null distribution. Lastly, we assessed the importance of individual SGBs in predicting behavior using Python’s ‘scikit-learn’ (v1.6)^103^. Throughout the analyses, a *p* < 0.05 was considered statistically significant, except where otherwise indicated. All plots were created with ‘ggplot2’ (v3.4.1)^104^. All figure panels were assembled using Inkscape 1.2.

## Author Contributions

M.V.F.R., Conceptualization, Methodology, Investigation, Formal analysis, Visualization, Writing – original draft, Writing – review & editing; M.N.V., Conceptualization, Methodology, Writing – review & editing; M.J.S., Conceptualization, Methodology, Investigation, Writing – review & editing; A.H.M., Conceptualization, Methodology, Investigation, Formal analysis, Visualization, Writing – original draft, Writing – review & editing, Funding acquisition.

## Acknowledgments

Funding was provided by the National Institutes of Health (NIH) grants R35 GM138284 and R01 DK139214 to A.H.M. NIH had no role in study design, data collection, and interpretation, or the decision to submit the work for publication. We acknowledge Cornell’s Biotechnology Resource Center for the help with sequencing host transcriptomic and fecal metagenomic data. We also thank Sylvia Bayrakdarian for her help with the animal care and behavioral tests.

## Data availability

All the FASTQ sequence data and associated metadata have been deposited in NCBI’s Sequence Read Archive under accession no. PRJNA1271077. Code for all analyses conducted in this manuscript is available at https://github.com/CUMoellerLab/Real_etal_2025_Predator_Stress.

