## Supplemental Information for "The mouse gut microbiota responds to predator odor and predicts host behavior"

**Contains:**

Figures S1 to S8

Tables S1 to S7

Supplementary Data 1 to 5

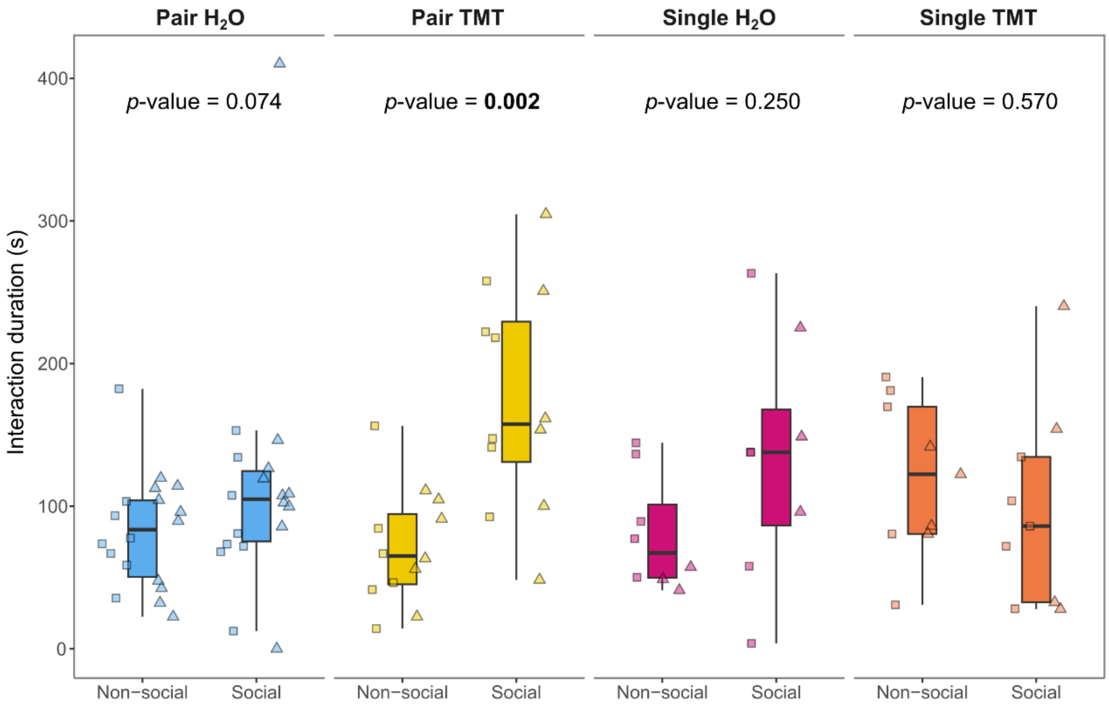


**Figure S1. Pair-housed mice exposed to predator odor displayed increased sociability.** Box plots show the amount of time that male (triangles) and female (squares) mice of different treatment groups spent interacting with a social stimulus (a stranger mouse) *vs*. a non-social stimulus (an empty cup) in the three-chamber test (3CT). Differences between the social and non-social interaction time within each treatment group were tested with a paired Wilcoxon test. Significant results ($p < 0.05$) are bolded.

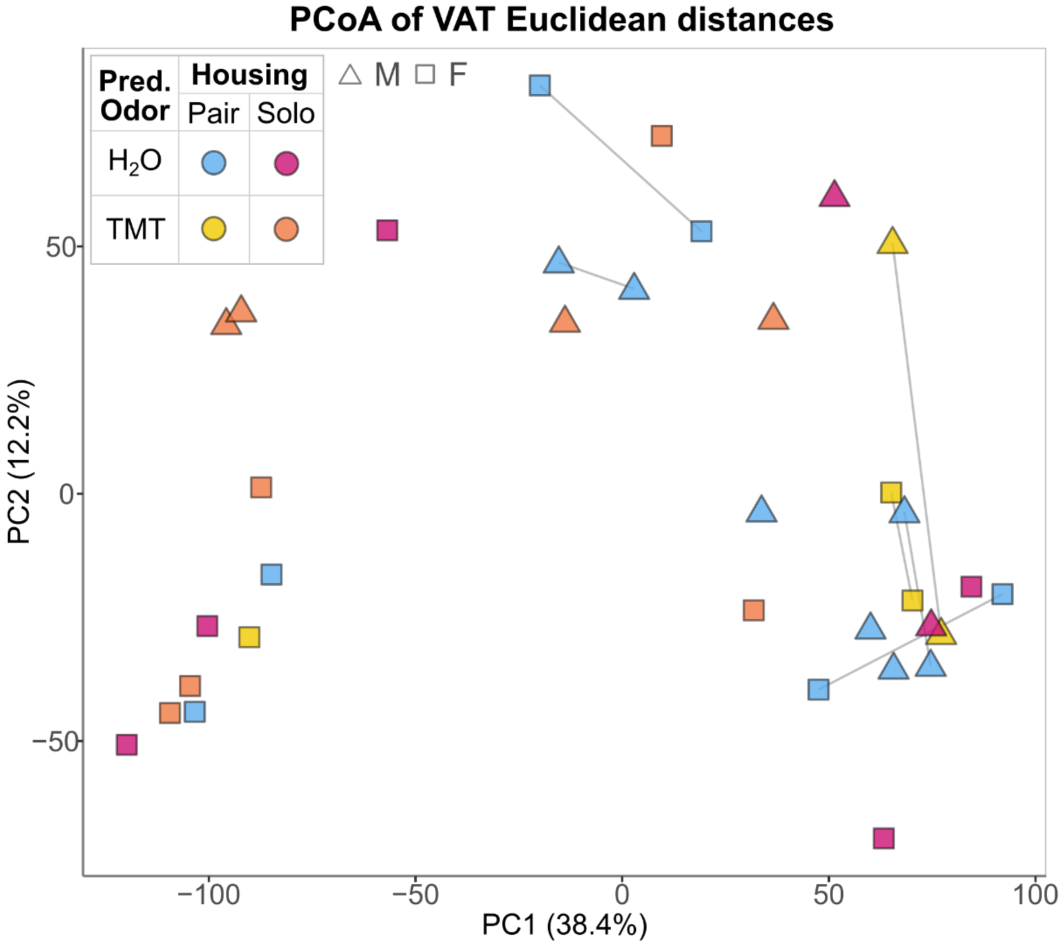


**Figure S2. Predator-odor exposure altered VAT gene expression.** Principal coordinate analysis (PCoA) shows the first two axes of the ordinated Euclidean distance matrix of the VAT transcriptome at the end of the experiment (D33). Points represent the transcriptome of male (triangles) and female (squares) mice, colored by treatment group: blue, Pair $H_2O$; yellow, Pair TMT; pink, Single $H_2O$; orange, Single TMT. A grey line connects pair-housed cage-mates. The percent variation explained by each axis is enclosed in parentheses.

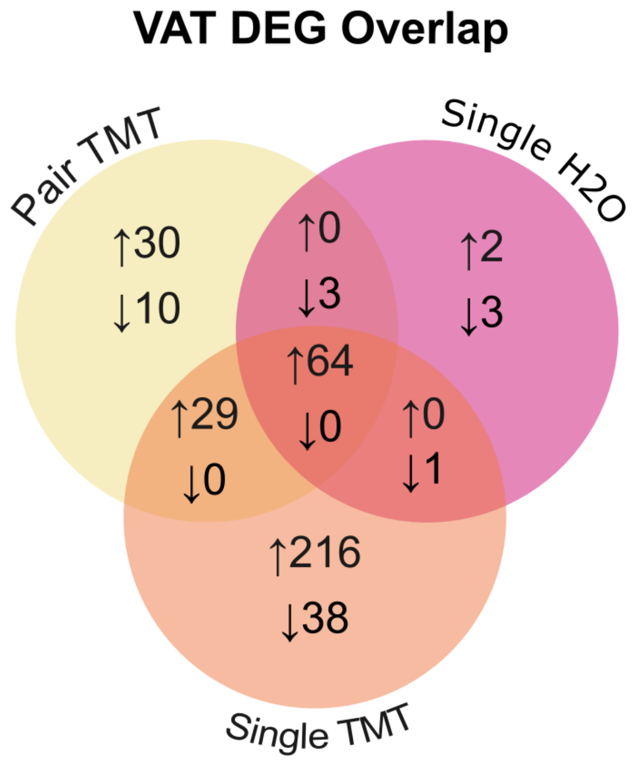


**Figure S3. Shared and unique DEGs among the stressor treatments.** Venn diagram shows the overlap in differentially expressed genes (DEGs) in the VAT of mice exposed to the stressors (Pair TMT, Single $H_2O$, and Single TMT) compared to unstressed controls (Pair $H_2O$). The upward arrow denotes up-regulated genes and the downward arrow denotes down-regulated genes.

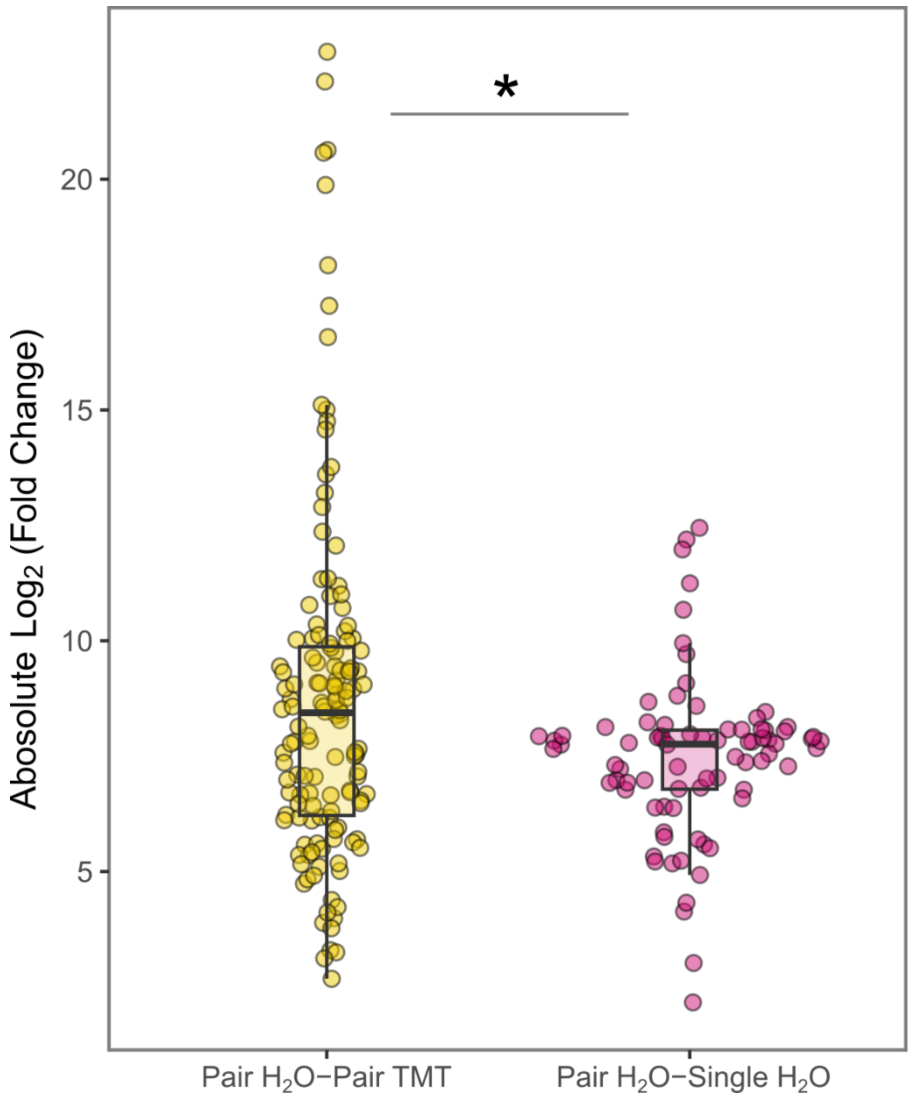


**Figure S4. Predator odor caused a larger mean fold-change in gene expression than did social isolation.** Box plot shows the absolute $log_2$(fold-change) in gene expression of DEGs in the VAT of pair-housed mice exposed to TMT (yellow) and single-housed mice exposed to $H_2O$ (pink), when compared to unstressed controls (Pair $H_2O$). The difference between the groups was tested with an unpaired Wilcoxon test (* = $p < 0.05$).

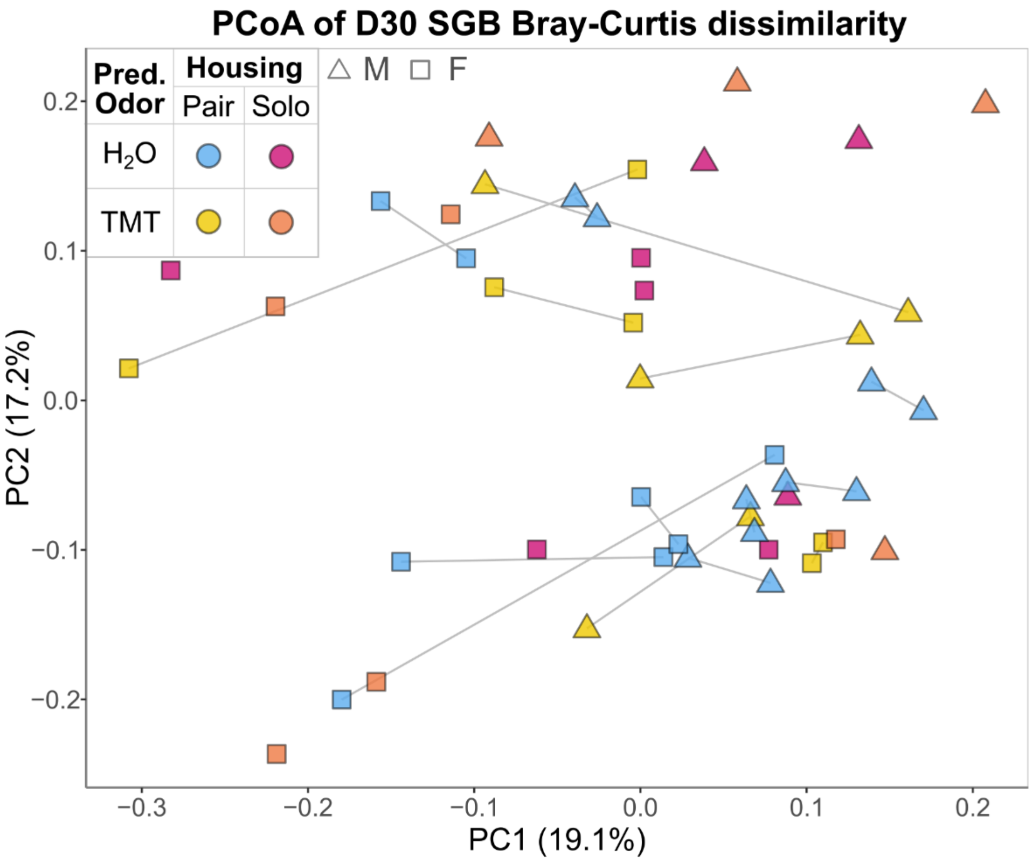


**Figure S5. Predator-odor exposure shifted the mean gut microbiota composition.** Principal coordinate analysis (PCoA) shows the first two axes of the ordinated Bray-Curtis dissimilarity matrix of species genome bins (SGBs) composition at the end of the experiment (D30). Points represent the species profiles of male (triangles) and female (squares) mice, colored by treatment group: blue, Pair H2O; yellow, Pair TMT; pink, Single H2O; orange, Single TMT. Pair-housed cage-mates are connected by a gray line. The percent variation in SGB composition explained by each axis is enclosed in parenthesis.

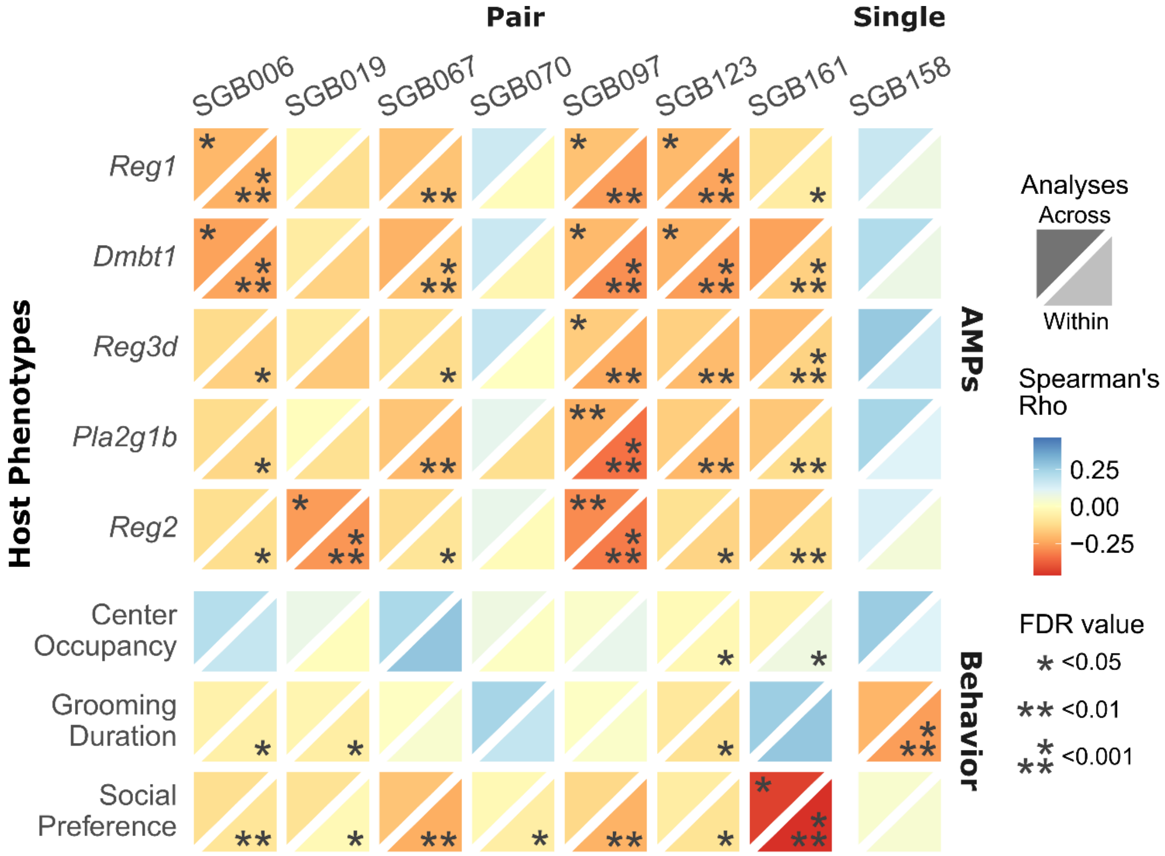


**Figure S6. Social isolation–responsive SGBs co-varied with host anti-microbial and behavioral responses.** Heatmap shows associations between the D22 relative abundances of SGBs that were consistently associated with social isolation (differentially abundant at both D22 and D30) and host anti-microbial (AMPs) and behavioral responses. The top facet represents correlations across treatment groups, and the bottom facet correlations within groups. Facets are colored according to Spearman's Rho and annotated with the correlation's FDR-adjusted *p*-value (* < 0.05; ** < 0.01; *** < 0.001).

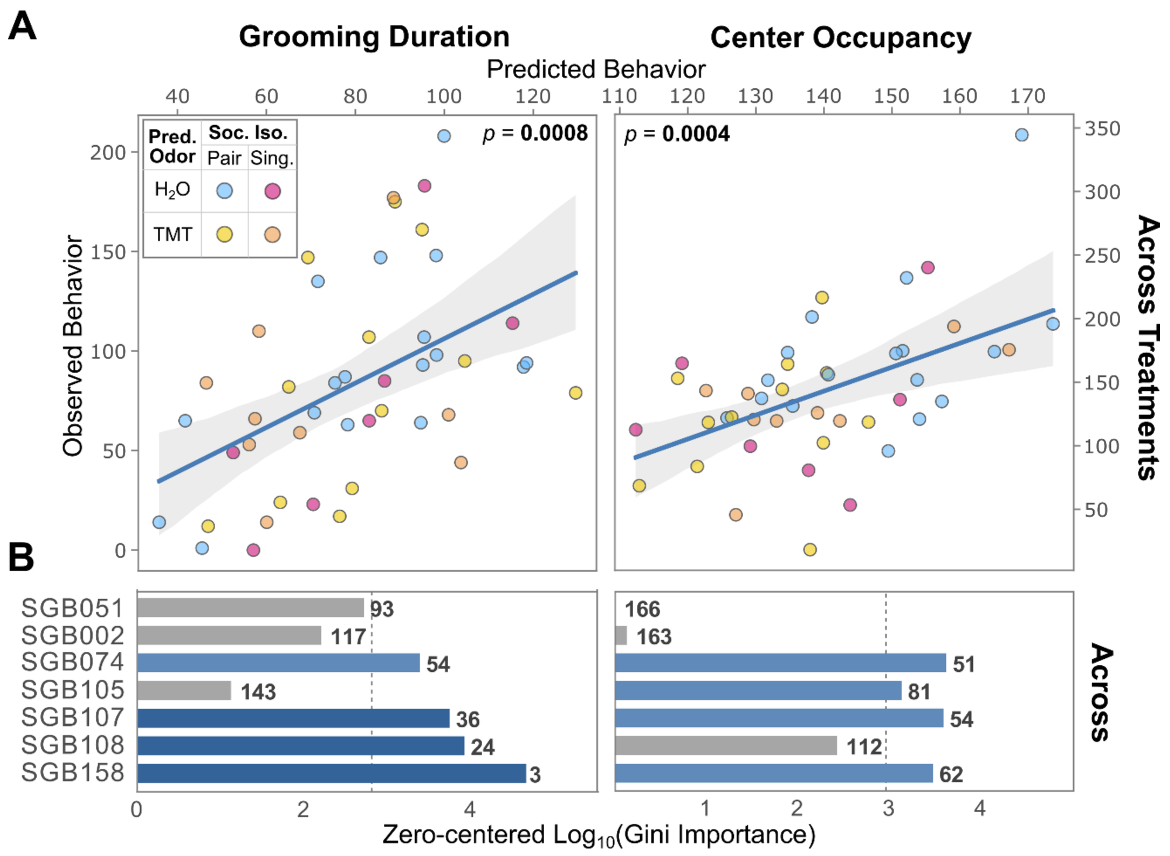


**Figure S7. Gut microbiota composition predicted host behavior measured a week later.** (**A**) Scatter plots show the relation between observed host behaviors – grooming (left column) and fearfulness (right column) – and the values predicted by a random forest model based on SGB relative abundances at D22 across treatment groups. Points are colored by treatment group: blue, Pair $H_2O$; yellow, Pair TMT; pink, Single $H_2O$; orange, Single TMT. The significance of the linear regression is shown in the upper inner corner, with significant values ($p < 0.05$) in bold. (**B**) The bar plots show the zero-centered $\log_{10}$(Gini Importance) of SGBs consistently associated with predator-odor exposure (differentially abundant at both D22 and D30) for predicting grooming behavior (left) and fearfulness (right) across treatment groups. The SGB importance rank order is annotated in bold. The bars are colored light blue if the SGB was among the top half most important features and dark blue if among the top quarter. The dashed line indicates the median importance value.

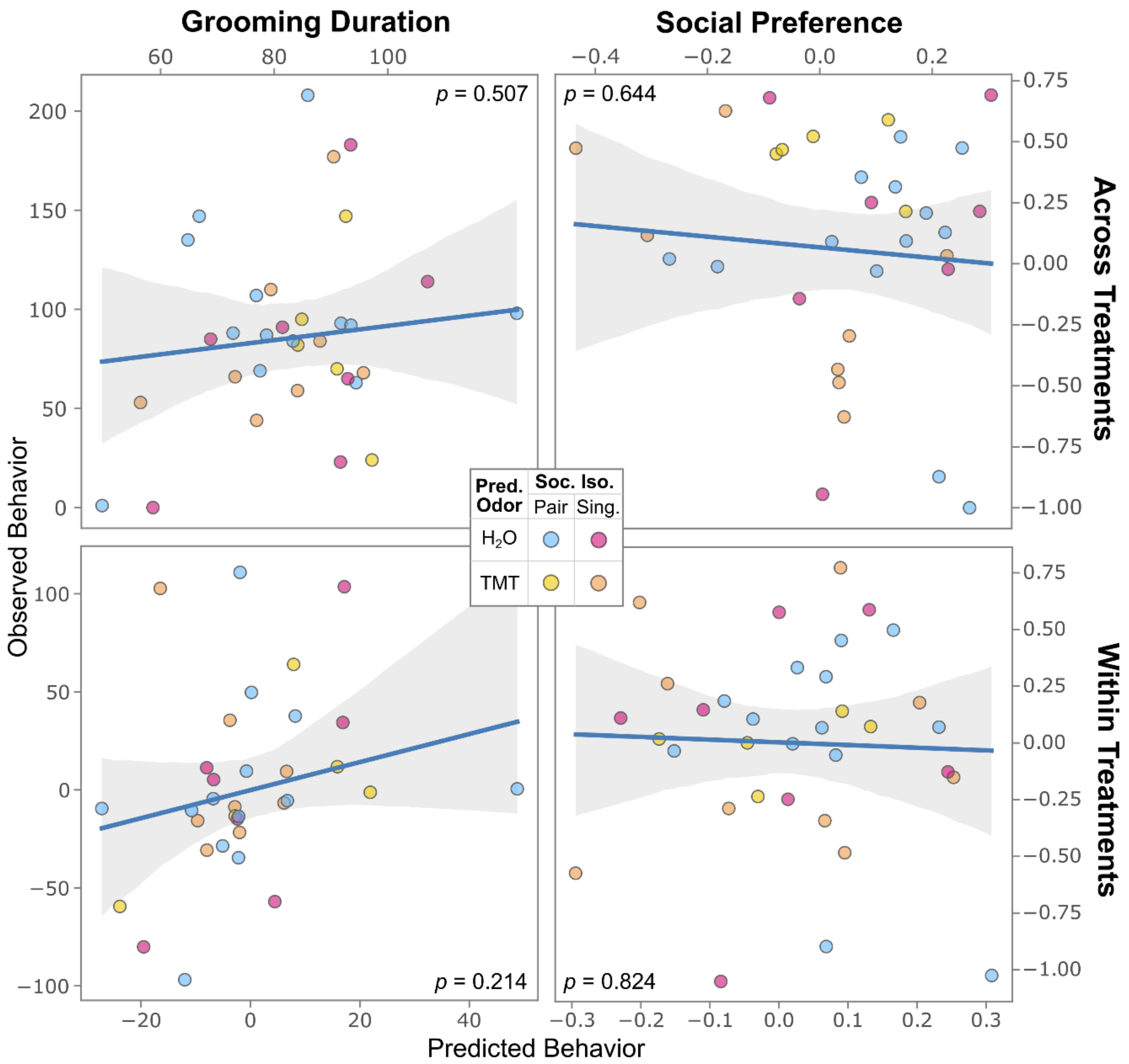


**Figure S8. VAT gene expression was not predictive of host behavior.** Scatter plots show the relation between observed grooming (left column) and social (right column) behavior and the values predicted by a random forest model based on VAT gene expression, both across (top row) and within treatment groups (bottom row; behavior residualized to account for the effect of stressor treatment). Points are colored by treatment group: blue, Pair $H_2O$; yellow, Pair TMT; pink, Single $H_2O$; orange, Single TMT. The significance of the linear regression is shown in the inner corner.

**Table S1.** Results from linear mixed-effects model (LMM) showing effects of sex, predator odor, social isolation, or their combination on mouse behavior. Significant differences ($p < 0.05$) are in bold.

| Behavior | Factor | Estimate | SE | DF | *t* | *p* |
|---|---|---|---|---|---|---|
| Center Occupancy | Sex | 0.008 | 0.027 | 40.682 | 0.298 | 0.767 |
| | Pair H2O – Pair TMT | -0.093 | 0.038 | 26.584 | -2.426 | **0.022** |
| | Pair H2O – Single H2O | -0.084 | 0.041 | 39.285 | -2.039 | **0.048** |
| | Pair H2O – Single TMT | -0.072 | 0.039 | 39.910 | -1.831 | 0.075 |
| Grooming Duration | Sex | 23.881 | 11.801 | 35.951 | 2.024 | 0.051 |
| | Pair H2O – Pair TMT | -26.043 | 20.615 | 41.058 | -1.263 | 0.214 |
| | Pair H2O – Single H2O | -44.327 | 20.299 | 41.979 | -2.184 | **0.035** |
| | Pair H2O – Single TMT | -39.629 | 19.204 | 41.936 | -2.064 | **0.045** |
| Social Preference | Sex | -0.105 | 0.126 | 40.395 | -0.830 | 0.411 |
| | Pair H2O – Pair TMT | 0.271 | 0.182 | 26.281 | 1.488 | 0.149 |
| | Pair H2O – Single H2O | 0.083 | 0.194 | 39.015 | 0.430 | 0.669 |
| | Pair H2O – Single TMT | -0.242 | 0.185 | 39.733 | -1.311 | 0.198 |
| Total Locomotion | Sex | 0.142 | 0.433 | 19.473 | 0.328 | 0.747 |
| | Pair H2O – Pair TMT | -0.755 | 0.647 | 20.452 | -1.167 | 0.257 |
| | Pair H2O – Single H2O | -0.364 | 0.666 | 32.941 | -0.547 | 0.588 |
| | Pair H2O – Single TMT | -0.186 | 0.635 | 32.229 | -0.294 | 0.771 |

**Table S2.** Results from pairwise LMM showing the differences in behavior between the different stressor treatments. Significant differences after Benjamini-Hochberg correction (FDR < 0.05) are in bold.

| Behavior | Contrast | Estimate | SE | *t* | *p* | FDR |
|---|---|---|---|---|---|---|
| Center Occupancy | Pair TMT – Single H2O | -0.009 | 0.044 | -0.202 | 0.840 | 0.840 |
| | Pair TMT – Single TMT | -0.021 | 0.042 | -0.502 | 0.616 | 0.840 |
| | Single H2O – Single TMT | -0.012 | 0.044 | -0.274 | 0.784 | 0.840 |
| Grooming Duration | Pair TMT – Single H2O | 18.284 | 21.045 | 0.869 | 0.385 | 0.715 |
| | Pair TMT – Single TMT | 13.586 | 19.098 | 0.711 | 0.477 | 0.715 |
| | Single H2O – Single TMT | -4.698 | 18.213 | -0.258 | 0.796 | 0.796 |
| Social Preference | Pair TMT – Single H2O | 0.187 | 0.210 | 0.894 | 0.371 | 0.371 |
| | Pair TMT – Single TMT | 0.513 | 0.197 | 2.610 | 0.009 | **0.027** |
| | Single H2O – Single TMT | 0.326 | 0.204 | 1.594 | 0.111 | 0.166 |
| Total Locomotion | Pair TMT – Single H2O | -0.391 | 0.721 | -0.542 | 0.588 | 0.793 |
| | Pair TMT – Single TMT | -0.569 | 0.673 | -0.845 | 0.398 | 0.793 |
| | Single H2O – Single TMT | -0.178 | 0.678 | -0.262 | 0.793 | 0.793 |

**Table S3.** Results from PERMANOVA and PERMDISP analyses showing the effect of predator odor (TMT) and social isolation (Housing) on the Euclidean distances of the visceral adipose tissue (VAT) transcriptome. Significant stressor effects ($p$ < 0.05) are in bold.

| Test | Factor | DF | Sum of Squares | $R^2$ | *F* | *p* |
|---|---|---|---|---|---|---|
| PERMANOVA | Sex | 1 | 16444.484 | 0.036 | 1.666 | 0.099 |
| | Litter | 10 | 195428.974 | 0.430 | 1.980 | 0.002 |
| | Housing | 1 | 16593.089 | 0.037 | 1.681 | 0.097 |
| | TMT | 1 | 21563.061 | 0.047 | 2.185 | **0.035** |
| | Residual | 20 | 197377.341 | 0.434 | | |
| | Total | 33 | 454424.543 | 1.000 | | |
| PERMDISP | Housing | 1 | 145.910 | | 0.242 | 0.647 |
| | Housing:Residuals | 32 | 19273.247 | | | |
| | TMT | 1 | 264.474 | | 0.498 | 0.505 |
| | TMT:Residuals | 32 | 16998.042 | | | |

**Table S4.** Significant results (FDR-adjusted $p < 0.05$) from the biological pathway enrichment analyses of differentially expressed genes (DEGs) in the VAT of TMT-exposed (Pair TMT) and single-housed mice (Single $H_2O$) when compared to unstressed controls (Pair $H_2O$). The gene ontology (GO) containing genes involved in antimicrobial humoral immune response mediated by antimicrobial peptides (AMPs) is in bold.

| Comparison | GO ID | Description | Gene Ratio | Bg Ratio | *p* | FDR |
|---|---|---|---|---|---|---|
| Pair $H_2O$ - Pair TMT | GO:0031016 | pancreas development | 7/127 | 75/17475 | 1.194E-06 | 0.002 |
| | GO:0031638 | zymogen activation | 6/127 | 66/17475 | 8.321E-06 | 0.007 |
| | **GO:0061844** | **antimicrobial humoral immune response mediated by antimicrobial peptide** | **6/127** | **75/17475** | **1.748E-05** | **0.007** |
| | GO:0031018 | endocrine pancreas development | 5/127 | 45/17475 | 1.813E-05 | 0.007 |
| | GO:0016485 | protein processing | 9/127 | 249/17475 | 8.619E-05 | 0.028 |
| | GO:0043588 | skin development | 9/127 | 266/17475 | 1.422E-04 | 0.036 |
| | GO:0007586 | digestion | 6/127 | 113/17475 | 1.752E-04 | 0.036 |
| | GO:0035270 | endocrine system development | 6/127 | 113/17475 | 1.752E-04 | 0.036 |
| | GO:0019730 | antimicrobial humoral response | 6/127 | 116/17475 | 2.022E-04 | 0.037 |

| | | | | | | |
|---|---|---|---|---|---|---|
| Pair $H_2O$ - Single $H_2O$ | GO:0007586 | digestion | 7/75 | 113/17475 | 5.414E-07 | 0.001 |
| | GO:0007631 | feeding behavior | 6/75 | 109/17475 | 7.279E-06 | 0.004 |
| | GO:0031016 | pancreas development | 5/75 | 75/17475 | 1.737E-05 | 0.004 |
| | **GO:0061844** | **antimicrobial humoral immune response mediated by antimicrobial peptide** | **5/75** | **75/17475** | **1.737E-05** | **0.004** |
| | GO:0019730 | antimicrobial humoral response | 5/75 | 116/17475 | 1.408E-04 | 0.029 |
| | GO:0031638 | zymogen activation | 4/75 | 66/17475 | 1.844E-04 | 0.031 |

**Table S5.** Results from PERMANOVA and PERMDISP analyses showing the effect of predator odor (TMT) and social isolation (Housing) on the Bray-Curtis dissimilarities in species-level genome bins (SGB) composition. A Benjamini–Hochberg correction was applied to the *p*-values to account for the repeated hypothesis testing across four different timepoints. Significant stressor effects (FDR < 0.05) are in bold.

| Test | Time point | Factor | DF | Sum of Squares | $R^2$ | F | *p* | FDR |
|---|---|---|---|---|---|---|---|---|
| PERMANOVA | D0 | Sex | 1 | 0.132 | 0.025 | 2.490 | 0.022 | 0.022 |
| | | Litter | 10 | 2.870 | 0.544 | 5.397 | 0.001 | 0.001 |
| | | Housing | 1 | 0.074 | 0.014 | 1.393 | 0.180 | 0.277 |
| | | TMT | 1 | 0.085 | 0.016 | 1.604 | 0.122 | 0.163 |
| | | Cage_ID | 7 | 0.732 | 0.139 | 1.966 | 0.002 | 0.004 |
| | | Residual | 26 | 1.383 | 0.262 | | | |
| | | Total | 46 | 5.276 | 1.000 | | | |
| | D15 | Sex | 1 | 0.193 | 0.057 | 6.488 | 0.001 | 0.001 |
| | | Litter | 10 | 1.876 | 0.557 | 6.308 | 0.001 | 0.001 |
| | | Housing | 1 | 0.025 | 0.007 | 0.844 | 0.590 | 0.590 |
| | | TMT | 1 | 0.035 | 0.011 | 1.192 | 0.285 | 0.285 |
| | | Cage_ID | 17 | 0.909 | 0.270 | 1.798 | 0.004 | 0.005 |
| | | Residual | 11 | 0.327 | 0.097 | | | |
| | | Total | 41 | 3.365 | 1.000 | | | |
| | D22 | Sex | 1 | 0.249 | 0.062 | 5.418 | 0.001 | 0.001 |
| | | Litter | 10 | 1.853 | 0.464 | 4.032 | 0.001 | 0.001 |
| | | Housing | 1 | 0.117 | 0.029 | 2.546 | 0.002 | **0.008** |
| | | TMT | 1 | 0.073 | 0.018 | 1.581 | 0.106 | 0.163 |
| | | Cage_ID | 17 | 1.062 | 0.266 | 1.360 | 0.010 | 0.010 |
| | | Residual | 14 | 0.643 | 0.161 | | | |
| | | Total | 44 | 3.998 | 1.000 | | | |
| | D30 | Sex | 1 | 0.266 | 0.071 | 8.022 | 0.001 | 0.001 |
| | | Litter | 10 | 1.763 | 0.474 | 5.320 | 0.001 | 0.001 |
| | | Housing | 1 | 0.042 | 0.011 | 1.282 | 0.208 | 0.277 |
| | | TMT | 1 | 0.097 | 0.026 | 2.928 | 0.002 | **0.008** |
| | | Cage_ID | 18 | 1.053 | 0.283 | 1.765 | 0.002 | 0.004 |
| | | Residual | 15 | 0.497 | 0.134 | | | |
| | | Total | 46 | 3.718 | 1.000 | | | |
| PERMDISP | D0 | Housing | 1 | 0.009 | | 1.354 | 0.275 | 0.435 |
| | | Housing:Residuals | 45 | 0.297 | | | | |
| | | TMT | 1 | 0.005 | | 0.659 | 0.404 | 0.791 |
| | | TMT:Residuals | 45 | 0.355 | | | | |
| | D15 | Housing | 1 | 0.005 | | 0.876 | 0.356 | 0.435 |

| | | | | | | | | |
|---|---|---|---|---|---|---|---|---|
| | | Housing:Residuals | 40 | 0.234 | | | | |
| | | TMT | 1 | 0.002 | | 0.268 | 0.587 | 0.791 |
| | | TMT:Residuals | 40 | 0.242 | | | | |
| | D22 | Housing | 1 | 0.002 | | 0.620 | 0.435 | 0.435 |
| | | Housing:Residuals | 43 | 0.155 | | | | |
| | | TMT | 1 | 0.001 | | 0.275 | 0.593 | 0.791 |
| | | TMT:Residuals | 43 | 0.175 | | | | |
| | D30 | Housing | 1 | 0.020 | | 5.754 | 0.019 | 0.076 |
| | | Housing:Residuals | 45 | 0.160 | | | | |
| | | TMT | 1 | 0.000 | | 0.006 | 0.936 | 0.936 |
| | | TMT:Residuals | 45 | 0.195 | | | | |

**Table S6.** Results from random forest models testing the predictive power of SGB abundances and VAT transcriptome on host behavior, both across and within groups (behavior residualized to account for the effect of stressor treatment). Significant results after FDR correction (FDR < 0.05) are in bold.

| Predictor | Analysis | Timepoint | Variable | $R^2$ | *p* | FDR |
|---|---|---|---|---|---|---|
| **Gut Microbiota** | Across | D0 | Center occupancy | -0.482 | 0.495 | 0.495 |
| | | | Grooming duration | -0.020 | 0.029 | **0.039** |
| | | | Social preference | -0.130 | 0.100 | 0.200 |
| | | D15 | Center occupancy | -0.169 | 0.083 | 0.166 |
| | | | Grooming duration | -0.369 | 0.412 | 0.412 |
| | | | Social preference | -0.327 | 0.323 | 0.430 |
| | | D22 | Center occupancy | 0.156 | 0.004 | **0.016** |
| | | | Grooming duration | 0.162 | 0.002 | **0.008** |
| | | | Social preference | -0.603 | 0.739 | 0.739 |
| | | D30 | Center occupancy | -0.278 | 0.215 | 0.286 |
| | | | Grooming duration | 0.065 | 0.009 | **0.018** |
| | | | Social preference | 0.151 | 0.002 | **0.008** |
| | Within | D0 | Center occupancy | -0.523 | 0.572 | 0.623 |
| | | | Grooming duration | -0.011 | 0.033 | 0.066 |
| | | | Social preference | -0.228 | 0.171 | 0.262 |
| | | D15 | Center occupancy | -0.256 | 0.159 | 0.474 |
| | | | Grooming duration | -0.587 | 0.748 | 0.748 |
| | | | Social preference | -0.243 | 0.197 | 0.262 |

| | | | | | | |
|---|---|---|---|---|---|---|
| | | D22 | Center occupancy | -0.290 | 0.237 | 0.474 |
| | | | Grooming duration | -0.165 | 0.139 | 0.185 |
| | | | Social preference | -0.675 | 0.799 | 0.799 |
| | | D30 | Center occupancy | -0.545 | 0.623 | 0.623 |
| | | | Grooming duration | 0.021 | 0.016 | 0.064 |
| | | | Social preference | 0.102 | 0.001 | **0.004** |
| **VAT Transcriptome** | Across | D33 | Center occupancy | -0.650 | 0.673 | 0.673 |
| | | | Grooming duration | -0.071 | 0.033 | **0.033** |
| | | | Social preference | -0.584 | 0.567 | 0.567 |
| | Within | D33 | Center occupancy | -0.899 | 0.835 | 0.835 |
| | | | Grooming duration | -0.467 | 0.454 | 0.454 |
| | | | Social preference | -0.902 | 0.852 | 0.852 |

**Table S7.** Summary results of the predictive power (Gini Importance) of SGBs that were consistently associated (DA at both D22 and D30) with predator-odor exposure or social isolation in regard to host behavior. SGBs that ranked among the top half most important features are in **bold**, and those in the top quarter are also **<u>underlined</u>**.

| Stressor | Analysis | Timepoint | Variable | SGB ID | Gini Importance | Rank |
|---|---|---|---|---|---|---|
| Predator Odor | Across | D22 | Center occupancy | SGB074 | 4.11E-03 | **51** |
| | | | | SGB107 | 3.82E-03 | **54** |
| | | | | SGB158 | 2.94E-03 | **62** |
| | | | | SGB105 | 1.34E-03 | **81** |
| | | | | SGB108 | 2.62E-04 | 112 |
| | | | | SGB002 | 3.42E-07 | 163 |
| | | | | SGB051 | 0.00E+00 | 166 |
| | | | Grooming duration | SGB158 | 4.49E-02 | **<u>3</u>** |
| | | | | SGB108 | 8.23E-03 | **<u>24</u>** |
| | | | | SGB107 | 5.45E-03 | **<u>36</u>** |
| | | | | SGB074 | 2.39E-03 | **54** |
| | | | | SGB051 | 5.08E-04 | 93 |
| | | | | SGB002 | 1.55E-04 | 117 |
| | | | | SGB105 | 1.20E-05 | 143 |
| | | | Social preference | SGB051 | 4.52E-03 | **<u>38</u>** |
| | | | | SGB158 | 4.33E-03 | **<u>40</u>** |
| | | | | SGB108 | 1.83E-03 | **55** |
| | | | | SGB107 | 1.30E-03 | **68** |
| | | | | SGB074 | 5.18E-04 | 97 |
| | | | | SGB105 | 3.47E-04 | 103 |
| | | | | SGB002 | 2.11E-04 | 112 |
| | | D30 | Center occupancy | SGB158 | 2.39E-02 | **<u>10</u>** |
| | | | | SGB002 | 3.93E-03 | **44** |
| | | | | SGB074 | 2.37E-03 | **62** |
| | | | | SGB108 | 1.85E-03 | **68** |
| | | | | SGB105 | 4.32E-04 | 102 |
| | | | | SGB051 | 1.64E-05 | 145 |
| | | | | SGB107 | 0.00E+00 | 166 |
| | | | Grooming duration | SGB051 | 6.28E-03 | **<u>36</u>** |
| | | | | SGB107 | 1.92E-03 | **64** |
| | | | | SGB158 | 1.40E-04 | 129 |
| | | | | SGB074 | 1.07E-04 | 132 |
| | | | | SGB108 | 2.01E-05 | 148 |
| | | | | SGB105 | 9.84E-06 | 151 |

| | | | | | | |
|---|---|---|---|---|---|---|
| | | | | SGB002 | 2.37E-07 | 165 |
| | | | Social preference | SGB074 | 1.68E-02 | <u>**19**</u> |
| | | | | SGB051 | 9.33E-03 | <u>**27**</u> |
| | | | | SGB105 | 4.92E-03 | <u>**42**</u> |
| | | | | SGB002 | 3.74E-03 | **46** |
| | | | | SGB158 | 1.69E-03 | **78** |
| | | | | SGB108 | 1.09E-04 | 129 |
| | | | | SGB107 | 1.29E-05 | 151 |
| | Within | D22 | Center occupancy | SGB158 | 6.02E-02 | <u>**2**</u> |
| | | | | SGB051 | 7.65E-03 | <u>**33**</u> |
| | | | | SGB107 | 6.59E-03 | <u>**34**</u> |
| | | | | SGB105 | 1.90E-03 | **72** |
| | | | | SGB074 | 1.07E-04 | 128 |
| | | | | SGB108 | 1.21E-05 | 148 |
| | | | | SGB002 | 0.00E+00 | 166 |
| | | | Grooming duration | SGB158 | 2.73E-02 | <u>**8**</u> |
| | | | | SGB108 | 7.55E-03 | <u>**30**</u> |
| | | | | SGB074 | 2.99E-03 | **50** |
| | | | | SGB105 | 1.08E-03 | **77** |
| | | | | SGB002 | 8.87E-04 | **80** |
| | | | | SGB107 | 6.77E-04 | **86** |
| | | | | SGB051 | 1.35E-04 | 117 |
| | | | Social preference | SGB002 | 1.65E-03 | **68** |
| | | | | SGB107 | 1.01E-03 | **81** |
| | | | | SGB074 | 2.17E-04 | 120 |
| | | | | SGB051 | 1.00E-04 | 131 |
| | | | | SGB158 | 4.31E-05 | 138 |
| | | | | SGB105 | 5.05E-06 | 152 |
| | | | | SGB108 | 0.00E+00 | 167 |
| | | D30 | Center occupancy | SGB074 | 1.20E-02 | <u>**22**</u> |
| | | | | SGB002 | 7.91E-03 | <u>**39**</u> |
| | | | | SGB107 | 1.04E-03 | 91 |
| | | | | SGB158 | 2.87E-04 | 118 |
| | | | | SGB051 | 1.90E-04 | 120 |
| | | | | SGB108 | 1.40E-04 | 123 |
| | | | | SGB105 | 0.00E+00 | 171 |
| | | | Grooming duration | SGB107 | 2.74E-03 | **60** |
| | | | | SGB158 | 7.97E-04 | **83** |
| | | | | SGB051 | 6.47E-04 | 89 |
| | | | | SGB074 | 6.12E-04 | 90 |
| | | | | SGB108 | 3.58E-04 | 103 |
| | | | | SGB002 | 1.72E-05 | 148 |

| | | | | | | |
|---|---|---|---|---|---|---|
| | | | | SGB105 | 0.00E+00 | 170 |
| | | | Social preference | SGB002 | 1.80E-02 | <u>**20**</u> |
| | | | | SGB105 | 7.17E-03 | <u>**39**</u> |
| | | | | SGB107 | 3.04E-03 | **63** |
| | | | | SGB158 | 2.33E-03 | **71** |
| | | | | SGB074 | 2.77E-04 | 116 |
| | | | | SGB051 | 0.00E+00 | 167 |
| | | | | SGB108 | 0.00E+00 | 169 |
| Social Isolation | Across | D22 | Center occupancy | SGB006 | 3.08E-02 | <u>**5**</u> |
| | | | | SGB070 | 9.52E-03 | <u>**30**</u> |
| | | | | SGB123 | 7.20E-03 | <u>**36**</u> |
| | | | | SGB067 | 5.76E-03 | **44** |
| | | | | SGB158 | 2.94E-03 | **62** |
| | | | | SGB161 | 1.79E-03 | **72** |
| | | | | SGB019 | 8.51E-04 | 91 |
| | | | | SGB097 | 7.50E-04 | 96 |
| | | | Grooming duration | SGB158 | 4.49E-02 | <u>**3**</u> |
| | | | | SGB070 | 5.87E-03 | <u>**32**</u> |
| | | | | SGB019 | 1.22E-03 | **70** |
| | | | | SGB097 | 7.79E-04 | **81** |
| | | | | SGB067 | 6.80E-04 | **85** |
| | | | | SGB161 | 6.07E-04 | 87 |
| | | | | SGB123 | 3.89E-04 | 100 |
| | | | | SGB006 | 0.00E+00 | 163 |
| | | | Social preference | SGB161 | 1.77E-02 | <u>**16**</u> |
| | | | | SGB158 | 4.33E-03 | **40** |
| | | | | SGB070 | 4.27E-03 | **41** |
| | | | | SGB123 | 6.71E-04 | 87 |
| | | | | SGB006 | 2.81E-04 | 108 |
| | | | | SGB019 | 1.10E-04 | 120 |
| | | | | SGB067 | 8.17E-05 | 125 |
| | | | | SGB097 | 1.03E-08 | 165 |
| | | D30 | Center occupancy | SGB158 | 2.39E-02 | <u>**10**</u> |
| | | | | SGB070 | 5.99E-03 | <u>**34**</u> |
| | | | | SGB097 | 3.23E-03 | **48** |
| | | | | SGB006 | 1.54E-03 | **73** |
| | | | | SGB123 | 7.85E-05 | 131 |
| | | | | SGB161 | 4.74E-05 | 137 |
| | | | | SGB019 | 4.34E-06 | 157 |
| | | | | SGB067 | 3.01E-07 | 163 |
| | | | Grooming duration | SGB161 | 2.22E-02 | <u>**10**</u> |
| | | | | SGB006 | 1.60E-02 | <u>**16**</u> |

| | | | | | | |
|---|---|---|---|---|---|---|
| | | | | SGB019 | 2.59E-03 | **57** |
| | | | | SGB067 | 2.29E-03 | **58** |
| | | | | SGB070 | 4.25E-04 | 103 |
| | | | | SGB097 | 1.44E-04 | 128 |
| | | | | SGB158 | 1.40E-04 | 129 |
| | | | | SGB123 | 0.00E+00 | 167 |
| | | | Social preference | SGB067 | 1.94E-02 | **<u>15</u>** |
| | | | | SGB161 | 3.52E-03 | **48** |
| | | | | SGB070 | 2.27E-03 | **65** |
| | | | | SGB158 | 1.69E-03 | **78** |
| | | | | SGB019 | 7.20E-04 | 90 |
| | | | | SGB123 | 1.67E-04 | 126 |
| | | | | SGB097 | 2.17E-05 | 146 |
| | | | | SGB006 | 0.00E+00 | 169 |
| | Within | D22 | Center occupancy | SGB158 | 6.02E-02 | **<u>2</u>** |
| | | | | SGB019 | 1.89E-02 | **<u>13</u>** |
| | | | | SGB123 | 2.66E-03 | **62** |
| | | | | SGB161 | 6.79E-05 | 134 |
| | | | | SGB097 | 5.02E-07 | 160 |
| | | | | SGB070 | 1.50E-07 | 163 |
| | | | | SGB006 | 0.00E+00 | 167 |
| | | | | SGB067 | 0.00E+00 | 169 |
| | | | Grooming duration | SGB158 | 2.73E-02 | **<u>8</u>** |
| | | | | SGB067 | 4.30E-04 | 91 |
| | | | | SGB006 | 4.22E-04 | 92 |
| | | | | SGB123 | 1.80E-04 | 109 |
| | | | | SGB161 | 9.14E-05 | 126 |
| | | | | SGB070 | 7.09E-05 | 132 |
| | | | | SGB019 | 7.00E-06 | 155 |
| | | | | SGB097 | 0.00E+00 | 167 |
| | | | Social preference | SGB006 | 5.17E-03 | **47** |
| | | | | SGB161 | 4.05E-04 | 103 |
| | | | | SGB070 | 2.24E-04 | 119 |
| | | | | SGB158 | 4.31E-05 | 138 |
| | | | | SGB097 | 1.85E-05 | 143 |
| | | | | SGB019 | 1.45E-05 | 146 |
| | | | | SGB123 | 0.00E+00 | 165 |
| | | | | SGB067 | 0.00E+00 | 166 |
| | | D30 | Center occupancy | SGB067 | 6.90E-04 | 102 |
| | | | | SGB123 | 6.25E-04 | 104 |
| | | | | SGB158 | 2.87E-04 | 118 |
| | | | | SGB097 | 1.81E-04 | 121 |

| | | | | | | |
|---|---|---|---|---|---|---|
| | | | | SGB161 | 1.22E-05 | 151 |
| | | | | SGB006 | 6.28E-06 | 153 |
| | | | | SGB070 | 4.95E-06 | 154 |
| | | | | SGB019 | 1.83E-08 | 162 |
| | | | Grooming duration | SGB161 | 1.96E-02 | **11** |
| | | | | SGB070 | 5.91E-03 | **33** |
| | | | | SGB019 | 4.35E-03 | **43** |
| | | | | SGB006 | 2.50E-03 | **62** |
| | | | | SGB123 | 2.33E-03 | **65** |
| | | | | SGB158 | 7.97E-04 | **83** |
| | | | | SGB067 | 1.63E-05 | 149 |
| | | | | SGB097 | 6.57E-07 | 162 |
| | | | Social preference | SGB097 | 2.72E-03 | **67** |
| | | | | SGB006 | 2.44E-03 | **70** |
| | | | | SGB158 | 2.33E-03 | **71** |
| | | | | SGB161 | 1.62E-03 | **80** |
| | | | | SGB067 | 5.19E-04 | 104 |
| | | | | SGB070 | 5.02E-05 | 142 |
| | | | | SGB123 | 2.18E-06 | 161 |
| | | | | SGB019 | 0.00E+00 | 170 |

**Supplementary Data 1 (separate file).** Behavioral assays data and metadata.

**Supplementary Data 2 (separate file).** Visceral adipose tissue (VAT) normalized gene counts data.

**Supplementary Data 3 (separate file).** Results from DESeq2 analyses showing the effect of predator odor (TMT) and social isolation (Housing) on VAT gene expression.

**Supplementary Data 4 (separate file).** Fecal metagenomic data, including A) sample metadata, B) GTDB taxonomic classification of the representative species-genome bins (SGBs) in our dataset, and C) unrarefied (raw) and D) rarefied abundance table of representative SGBs.

**Supplementary Data 5 (separate file).** Results from DESeq2 analyses showing the effect of predator odor (TMT) and social isolation (Housing) on SGB abundance at D22 (T4) and D30 (T5).
